# The USP12/46 deubiquitinases protect integrins from ESCRT-mediated lysosomal degradation

**DOI:** 10.1101/2024.05.14.594138

**Authors:** Kaikai Yu, Shiny S. Guo, Florian Bassermann, Reinhard Fässler, Guan M. Wang

## Abstract

The functions of integrins are tightly regulated via multiple mechanisms including trafficking and degradation. Integrins are repeatedly internalized, routed into the endosomal system and either degraded by the lysosome or recycled back to the plasma membrane. The ubiquitin system dictates whether internalized proteins are degraded or recycled. Here, we used a genetic screen and proximity-dependent biotin identification to identify deubiquitinase(s) that control integrin surface levels. We found that a ternary deubiquitinating complex, comprised of USP12 (or the homologous USP46), WDR48 and WDR20, stabilizes β1 integrin (Itgb1) by preventing ESCRT-mediated lysosomal degradation. Mechanistically, the USP12/46-WDR48-WDR20 complex removes ubiquitin from the cytoplasmic tail of internalized Itgb1 in early endosomes, which in turn prevents ESCRT-mediated sorting and Itgb1 degradation.

## Introduction

Integrins are α/β heterodimers that mediate cell adhesion between cells and to the extracellular matrix (ECM) proteins (Hynes, 2002). The function of integrins is tightly regulated, on one hand by changing the conformational state that turns ligand binding on and off (Calderwood *et al*, 2013; Moser *et al*, 2009), and on the other hand by adjusting surface location and levels through an endosomal sorting process that dictates whether the integrins are recycled back to the cell surface or delivered to lysosomes for degradation (Moreno-Layseca *et al*, 2019). Considering that the approximate half-life of Itgb1-class integrins is 24-48 hours, their cell surface residence time 10 minutes and the recycling from and back to the plasma membrane around 20 minutes (Bottcher *et al*, 2012; Dozynkiewicz *et al*, 2012; Moreno-Layseca *et al*., 2019), it can be assumed that Itgb1-class integrins undergo numerous cycles of endocytosis and recycling during their lifespan before they are degraded in lysosomes.

The ubiquitin system is a labelling system that marks proteins for different proteolytic fates, such as integrins that are determined for lysosomal degradation. Integrins and other cell surface proteins designated for internalization are ubiquitin-tagged at lysine residues in their cytoplasmic tail (Clague *et al*, 2012). The removal of the ubiquitin tags by specific deubiquitinases (DUBs) in early endosomes directs proteins into the recycling pathway and back to the cell surface (Clague *et al*., 2012; Komander *et al*, 2009). Proteins, which retain the ubiquitin tag are recognized by the Endosomal Sorting Complex Required for Transport (ESCRT) complex, sequestered into microdomains and internalized as intraluminal vesicles (ILVs) leading to the formation of multivesicular bodies (MVBs), also known as late endosomes (Hanson & Cashikar, 2012). MVBs/late endosomes either mature into lysosomes in which transmembrane membrane proteins on ILVs are degraded by lysosomal proteases, or fuse with the plasma membrane which leads to the extracellular release of their cargo including the ILVs as exosomes (Huotari & Helenius, 2011; Saftig & Klumperman, 2009).

Previous studies have shown that the binding of α5β1 integrin to soluble fibronectin (FN) induces integrin cytoplasmic tail ubiquitination, internalization and the degradation of the integrin (Kharitidi *et al*, 2015; Lobert *et al*, 2010). It has also been demonstrated that internalized Itgb1 recruits the SNX17-retriever complex, which leads to the retrieval and recycling of integrins (Bottcher *et al*., 2012; McNally *et al*, 2017; Steinberg *et al*, 2012). Indeed, SNX17-binding-deficient Itgb1 tail mutants fail to recycle and are degraded in the lysosome. They can be rescued from degradation upon additionally substituting the α5β1 integrin tails lysines for non-ubiquitinatable arginines, which led to the hypothesis that SNX17 fulfils two functions: on one hand it recruits DUB(s) to deubiquitinate the Itgb1 tail (Bottcher *et al*., 2012), and on the other hand it recruits the retriever complex to retrieve and recycle integrins. USP9X has been identified to bind SNX17 and deubiquitinate centriolar satellite proteins required for ciliogenesis (Wang et al., 2019). Although integrin deubiquitination has not been investigated in this report, Kharitidi and colleagues showed in an independent study that USP9X can deubiquitinate the α5-subunit (Itga5) in cells upon treatment with soluble FN (Kharitidi *et al*., 2015). Importantly, however, tissues contain primarily FN that is crosslinked by the lysyl oxidase into an insoluble fibrillar network (Melamed *et al*, 2023) which, in contrast to soluble FN, cannot be internalized by integrins, raising the question whether USB9X also controls the steady state levels of unbound α5β1 integrins.

In the present paper, we designed unbiased genetic and biochemical screens aimed at identifying novel DUB(s) that stabilize Itgb1 levels at the cell surface at steady state. Our experiments revealed that the DUBs USP12 and USP46 complexed with WDR48 and WDR20 remove ubiquitin from the cytoplasmic tails of internalized Itgb1 and several other cells surface proteins including signalling proteins and solute transporters resulting in a decoupling from ESCRT-mediated degradation. The significance of our findings is discussed.

## Results

### The Itgb1 protein is stabilized by USP12 and USP46

As USP9X was shown to deubiquitinate Itga5 tails on endosomes following soluble FN stimulation (Kharitidi *et al*., 2015), we first investigated whether USP9X also deubiquitinates and stabilizes Itgb1 in cells cultured under steady state conditions. To this end, we cultured USP9X-depleted mouse fibroblasts, Hela, RPE-1 and MDA-MB-231 cells, respectively, either continuously in the presence of fetal bovine serum (10%, high concentration of soluble serum FN) or in the presence of serum replacement medium (which lacks soluble FN). The experiments revealed that depletion of USP9X in all cells analysed was either without effect or slightly increased rather than decreased Itga5 and Itgb1 levels on the surface which was measured by flow cytometry, and in lysates which was determined by Western blot (WB), irrespective whether exposed to medium containing or lacking FBS (Fig. EV1). These findings indicate that USP9X does not control integrin turnover under steady state culture conditions and suggests that unknown DUB(s) ensure retrieval of integrins.

These results prompted us to design an unbiased Crispr/Cas9-based genetic screen aimed at identifying DUBs that regulate the surface stability of Itgb1 in cells cultured with 10% FBS-containing medium at steady state (Fig. 1A). Specifically, we targeted 98 human DUBs genes by transducing the human Cas9-expressing haploid HAP1 cell line with pooled lentiviral guide RNA (gRNA) libraries (Paulmann *et al*, 2022). The transduced HAP1 cells were then expanded, fixed, immunostained for Itgb1 and sorted by flow cytometry to obtain the 5% cells with the lowest and the 5% cells with the highest Itgb1 surface levels. Next, we used next-generation sequencing (NGS) to identify the gRNA-targeted genes in the Itgb1^low^ and Itgb1^high^ cell populations, respectively. We identified *BAP1*, *USP7*, *OTUD6B* and *USP46* genes in the Itgb1^low^, and *PSMD14* and *USP14* in the Itgb1^high^ as potential regulators of Itgb1 cell surface levels (Fig. 1B).

**Figure 1.**
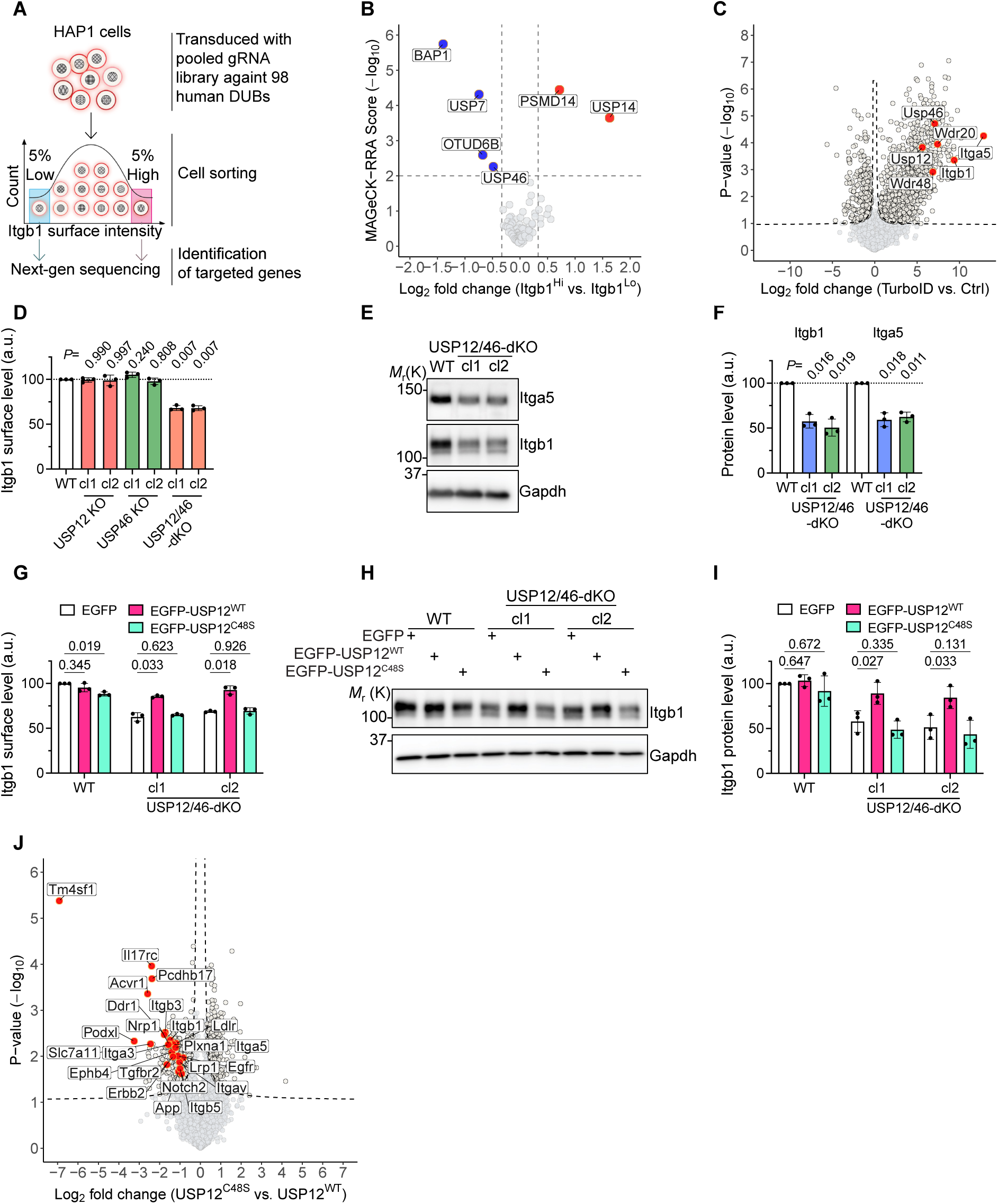
The USP12/46-WDRs complex stabilizes Itgb1 protein levels. **(A)** Schematic overview of the CRISPR screen for identifying DUBs regulating Itgb1 surface levels. Cas9-expressing HAP1 cells were transduced with pooled lentiviral guide RNA (gRNA) libraries targeting 98 DUBs from the human genome. After 2 weeks in culture, cells with the 5% lowest (Itgb1^Lo^) and the 5% highest (Itgb1^Hi^) Itgb1 surface levels were sorted by flow cytometry and gRNA-targeted genes were determined. **(B)** Volcano plot of the results from the CRISPR screen. The x-axis represents the log_2_ fold change (lfc) in the frequency of genes targeted between the Itgb1^Lo^ and Itgb1^Hi^ populations. The y-axis indicates the Robust Rank Aggregation (RRA) score determined by the MAGeCK algorithm (MAGeCK-RRA) (Li *et al*, 2014). Dots represent individual targeted genes, and those meeting the criteria of |lfc| > 0.33 and - log_10_(RRA) > 2 were considered significant. Genes significantly enriched in Itgb1^Lo^ cells are colored in blue and those enriched in Itgb1^Hi^ in red. **(C)** Volcano plot of the α5β1 integrin proximitome determined by label-free MS analysis in mouse kidney fibroblasts expressing miniTurbo-tagged Itag5 (TurboID) versus Itgb1-KO fibroblasts (Ctrl). *P*-values were determined using two-sided permuted *t*-test with 250 randomizations. The black dashed line indicates the significance cutoff (FDR:0.05, S0:0.1) estimated by the Perseus software. n=3 biological replicates. The red dots indicate the subunits of the α5β1 heterodimer and the components of the USP12/46-WDR48-WDR20 complex. **(D)** Itgb1 surface levels in WT and two independent clones (cl1 and cl2) of USP12-KO, USP46-KO and USP12/46-dKO fibroblasts determined by flow cytometry. Statistical analysis was carried out by RM one-way ANOVA with Dunnett’s multiple comparison test. Data are shown as Mean±SD, n=3 independent experiments. **(E, F)** WB (**E**) and densitometric quantification (**F**) of Itgb1 and Itga5 protein levels in WT and USP12/46-dKO fibroblasts. Gapdh served as loading control. Statistical analysis was carried out by RM one-way ANOVA with Dunnett’s multiple comparison test. Data are shown as Mean±SD, n=3 independent experiments. **(G-I)** Itgb1 surface levels determined by flow cytometry (**G**), Itgb1 protein levels in cell lysates determined by WB (**H**) and densitometric quantification (**I**) in WT and USP12/46-dKO fibroblasts stably expressing EGFP, EGFP-USP12^WT^ or EGFP-USP12^C48S^. Gapdh served as a loading control. Statistical analysis was carried out by RM two-way ANOVA with Dunnett’s multiple comparison test. Data are shown as Mean±SD, n=3 independent experiments. **(J)** Volcano plot of the cell surface proteome of USP12/46-dKO fibroblasts stably expressing EGFP-USP12^C48S^ versus EGFP-USP12^WT^ identified by label-free MS. *P*-values were determined using two-sided permuted *t*-test with 250 randomizations. The black dashed line indicates the significance cutoff (FDR:0.05, S0:0.1) estimated by the Perseus software. n=4 biological replicates. Arbitrarily selected cell surface receptors were highlighted in red.

To identify which of the DUBs identified in the Crispr/Cas9-based screen are present in the proximity of the Itgb1 tail in mouse fibroblasts, we determined the Itgb1 proximitome by combining the proximity-dependent biotin identification (BioID) assay in combination with mass spectrometry (MS)-based proteomics. First, we fused the miniTurbo (Branon *et al*, 2018) to the cytosolic tail of Itga5 which associates with Itgb1 whose tail integrity is required to bind interactors such as Kindlins and SNX17 (Bottcher *et al*., 2012; Fitzpatrick *et al*, 2014; Li *et al*, 2017). The Itga5-miniTurbo was retrovirally transduced into wild-type (WT) and Itgb1-KO fibroblasts. The newly synthesized Itga5 cannot heterodimerize in the absence of Itgb1 and is degraded in the endoplasmatic reticulum, which makes Itgb1-KO cells a perfect negative control. Next, we isolated biotinylated proteins from cell lysates with streptavidin-conjugated beads, performed MS and identified USP46, the paralog USP12, and the USP12- and USP46-binding and activating adaptor proteins WDR48 and WDR20 (Li *et al*, 2016; Zhu *et al*, 2019) (Fig. 1C). USP12 and USP46 share approximately 90% protein sequence similarity and conserved binding sites for WDR48 and WDR20 (Li *et al*., 2016; Zhu *et al*., 2019). The BAP1, USP7, OTUD6B and USP14 proteins were undetectable by MS. PSMD14 and USP9X were detected at comparably low levels in WT and Itgb1-KO fibroblasts, suggesting that these two proteins exhibit background binding, e.g. to the beads used in the experiment. Thus, the unbiased genetic screen as well as the proximitome point to the USP12/46-WDR48-WDR20 DUB complex (hereafter referred to as USP12/46-WDRs complex) as stabilizer of the Itgb1 surface levels.

To validate the results of our screens, we used Crispr/Cas9 technology to knockout (KO) the *USP12*, *USP46*, *WDR20*, and *WDR48* genes either individually or in combination in at least two different mouse fibroblast and human breast cancer MDA-MB-231 cell clones, respectively. Since antibodies against the USP12/46-WDRs complex are not available and several attempts to generate specific homemade polyclonal antisera were unsuccessful, we validated the KOs of the individual clones by genomic PCR followed by sequencing of the amplified genes (Fig. EV2). Flow cytometry analysis revealed that the expression levels of Itgb1 on fibroblasts carrying single KO of either USP12 or USP46 were comparable to those of wild-type fibroblasts (Fig. 1D), whereas the levels of Itgb1 on fibroblasts with the double knockout of USP12 and USP46 (USP12/46-dKO) were significantly reduced (Fig. 1D), suggesting that USP12 and USP46 compensate each other. WB of fibroblast lysates revealed that the levels of the 105 kDa Itgb1 band corresponding to the immature, ER-resident Itgb1 remained unaffected by the USP12/46-dKO, while the levels corresponding to the 125 kDa mature Itgb1 band were reduced in USP12/46-dKO cells (Fig. 1E, F), indicating that the destabilization occurs either in the secretory pathway, on the cell surface and/or in the endosomal system. Concomitantly with the decrease of the mature Itgb1 also the Itga5 levels were reduced in USP12/USP46-dKO lysates (Fig. 1E, F). In MDA-MB-231 cells, the dKO of USP12/46 also reduced the levels of Itgb1 at the cell surface and those of the 125 kDa mature Itgb1 in the whole cell lysate (Fig. EV3A-C).

The reduced Itgb1 levels in USP12/46-dKO fibroblasts were restored upon expression of either EGFP-tagged USP12^WT^ or Flag-tagged USP46^WT^ (Fig. 1G-I and Fig. EV3D-G). In contrast, re-expression of the catalytically inactive USP12^C48S^ or USP46^C44S^ mutants, in which the catalytic site cysteine was substituted for serine (Li *et al*., 2016; Yin *et al*, 2015), were unable to restore the Itgb1 levels indicating that USP12/46 require the DUB activity to stabilize the 125 kDa mature Itgb1 levels. Since USP12 and USP46 compensate each other, we used USP12 to delineate the DUB function in the following reconstitution experiments.

To assess whether USP12 regulates surface proteins other than Itgb1, we determined the cell surface proteome of USP12/46-dKO fibroblasts (Fig. 1J) and USP12/46-dKO MDA-MB-231 cells reconstituted with either USP12^WT^ or USP12^C48S^ (Fig. EV3H). To this end, we biotinylated cell surface proteins, precipitated the biotinylated proteins using streptavidin-conjugated beads and compared the abundance of the precipitated proteins by MS. We found that the levels of numerous surface receptors including integrins (Itgb3, Itgb5, Itga3 and Itgav), IL17rc, Pcdhb17, Acvr1, Ddr1, etc. were significantly decreased in USP12^C48S^ expressing fibroblasts (Fig. 1J). Decreased surface levels of integrins, FAT4, STEAP3, PLXNB3, FZD6, etc. were identified in USP12^C48S^ expressing MDA-MB-231 cells (Fig. EV3H). These results indicate that USP12 controls the levels of numerous surface proteins.

### Binding of USP12 to WDR48-WDR20 is essential to maintain Itgb1 surface levels

Previous studies have shown that the deubiquitinase activity of USP12 and USP46 requires the association with the adaptor proteins WDR48 or WDR20 and is further increased upon binding to both, WDR48 and WDR20 (Li *et al*., 2016; Zhu *et al*., 2019). In line with these findings, we observed that Crispr/Cas9-mediated KO of either WDR48 or WDR20 moderately decreased Itgb1 surface levels on independently generated fibroblast clones, whereas the dKO of WDR48 as well as WDR20 decreased Itgb1 surface levels to the same extent as in USP12/46-dKO or USP12^C48S^-expressing USP12/46-dKO fibroblasts (Fig. 2A). Furthermore, expression of either WDR48 or WDR20 alone in WDR48/20-dKO fibroblasts restored the levels of Itgb1 at the cell surface to a lesser extent than expression of the two WDR proteins together (Fig. 2B).

**Figure 2.**
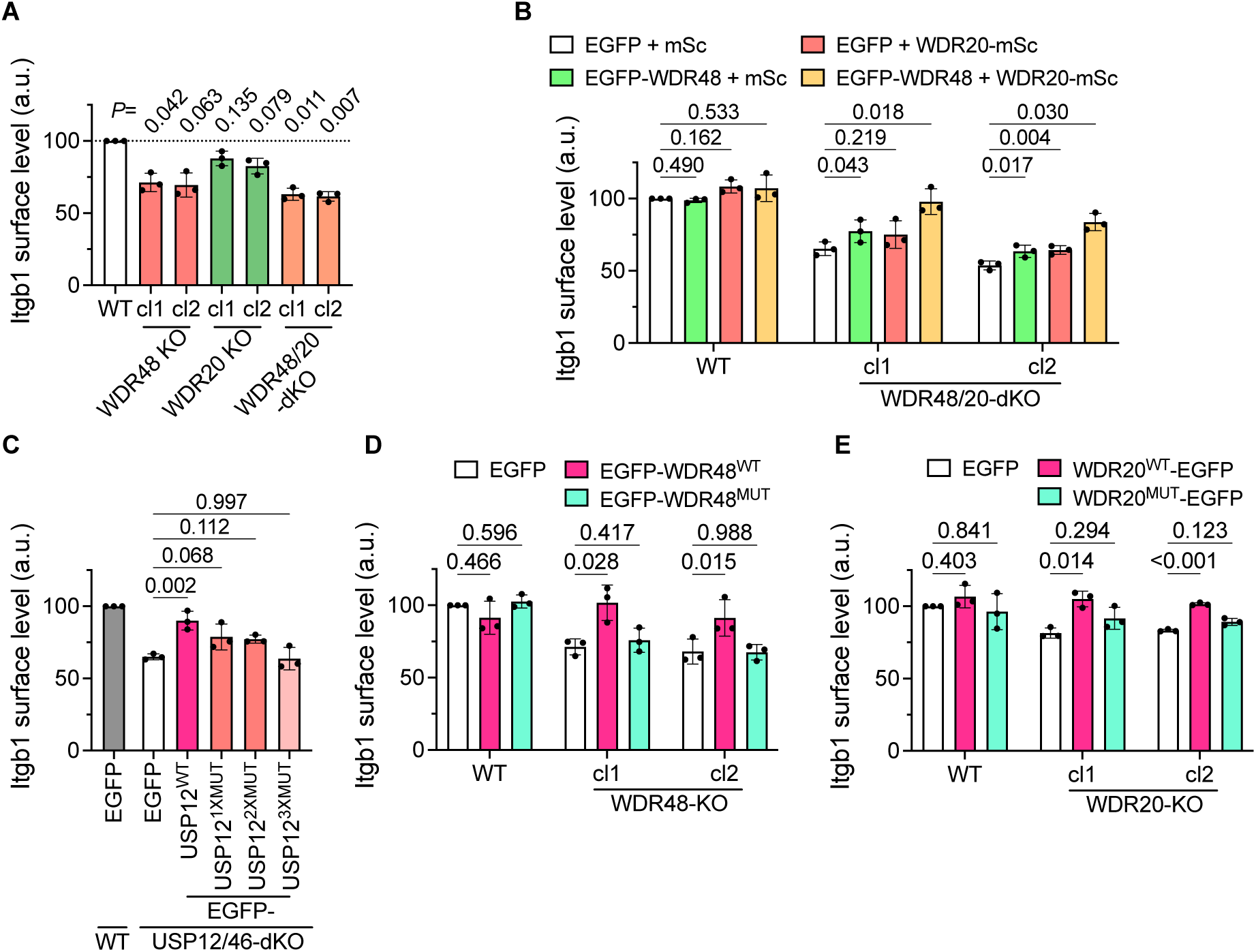
The integrity of the USP12/48-WDR20-WDR48 complex is required to stabilize Itgb1 protein levels. **(A)** Itgb1 surface levels in WT and WDR48-KO, WDR20-KO and WDR48/20-dKO fibroblasts determined by flow cytometry. Statistical analysis was carried out by RM one-way ANOVA with Dunnett’s multiple comparison test comparing with WT. Data are shown as Mean±SD, n=3 independent experiments. **(B)** Itgb1 surface levels in WT and WDR48/20-dKO fibroblasts transiently expressing EGFP, mScarlet, EGFP-WDR48 and/or WDR20-mScarlet determined by flow cytometry. Statistical analysis was carried out by RM two-way ANOVA with Dunnett’s multiple comparison test. Data are shown as Mean±SD, n=3 independent experiments. **(C)** Itgb1 surface levels in WT and USP12/46-dKO fibroblasts transiently expressing EGFP, EGFP-USP12^WT^, EGFP-USP12^1XMUT^ (E190K, deficient in binding to WDR48), EGFP-USP12^2XMUT^ (F287A, V279A, deficient in binding to WDR20) or EGFP-USP12^3XMUT^ (E190K, F287A, V279A, deficient in binding to WDR48 as well as WDR20) determined by flow cytometry. Statistical analysis was carried out by ordinary one-way ANOVA with Dunnett’s multiple comparison test. Data are shown as Mean±SD, n=3 independent experiments. **(D)** Itgb1 surface levels in WT and WDR48-KO fibroblasts transiently expressing EGFP-WDR48^WT^ or EGFP-WDR48^MUT^ (deficient in binding to USP12 as well as USP46) determined by flow cytometry. Statistical analysis was carried out by RM two-way ANOVA with Dunnett’s multiple comparison test. Data are shown as Mean±SD, n=3 independent experiments. **(E)** Itgb1 surface levels in WT and WDR20-KO fibroblasts transiently expressing WDR20^WT^-EGFP or WDR20^MUT^-EGFP (deficient in binding to USP12 as well as USP46) determined by flow cytometry. Statistical analysis was carried out by RM two-way ANOVA with Dunnett’s multiple comparison test. Data are shown as Mean±SD, n=3 independent experiments.

We also confirmed that the activity of USP12 depends on the direct interaction with WDR48 and WDR20 (Dharadhar *et al*, 2016; Li *et al*., 2016) by mutating the binding site in EGFP-tagged USP12 for WDR48 (USP12^1XMUT^, E190K) (Dharadhar *et al*., 2016), for WDR20 (USP12^2XMUT^, F287A, V279A) (Li *et al*., 2016) or for both, WDR48 and WDR20 (USP12^3XMUT^, E190K, F287A, V279A) (Fig. EV4A). Expression of USP12^WT^ in USP12/46-dKO fibroblasts rescued Itgb1 surface levels, whereas expression of USP12^1XMUT^ or USP12^2XMUT^ only partially rescued Itgb1 surface levels and expression of USP12^3XMUT^ did not increase Itgb1 levels beyond the levels of USP12/46-dKO fibroblasts expressing EGFP-only (Fig. 2C and Fig. EV4B). Also the expression of EGFP-tagged WDR48 mutant proteins (WDR48^MUT^, K214E/W256A/R272D) in WDR48-KO fibroblasts, and WDR20 mutant proteins (WDR20^MUT^, F262A/W306A) in WDR20-KO fibroblasts, both of which are unable to bind USP12/46 (Li *et al*., 2016; Yin *et al*., 2015), failed to normalize Itgb1 surface levels (Fig. 2D, E and Fig. EV4C-F). These findings indicate that the entire USP12/46-WDRs complex is required to stabilize Itgb1.

The WDR48 protein consists of an N-terminal β propeller domain followed by an ancillary domain (AD) and a C-terminal sumo-like domain (SLD), which is thought to recruit the substrate (in our case the ubiquitinated Itgb1 tail) to the USP-WDR48 complex (Li *et al*., 2016; Yin *et al*., 2015). However, expression of EGFP-tagged WDR48 protein lacking the SLD (WDR48^1-580^) or the SLD as well as the AD (WDR48^1-359^) domain in WDR48-KO fibroblasts restored Itgb1 surface levels to the same extent as expression of WDR48^WT^, indicating that neither the SLD nor the AD domains are required to control DUB-mediated Itgb1 surface levels (Fig. EV4G-I).

### SNX17 and the USP12/46-WDRs complex stabilize Itgb1 independently of each other

SNX17 binds Itgb1 on early endosomes and was shown to promote recycling of Itgb1 in an Itgb1-tail ubiquitination-dependent manner (Bottcher *et al*., 2012; Steinberg *et al*., 2012). Since dKO of USP12/46 reduced Itgb1 cell surface levels and total levels in cell lysates to a similar extent as loss of SNX17, we tested whether the deletion of the *Snx17* gene in USP12/46-dKO fibroblasts affects Itgb1 surface levels. The experiment revealed that the cell surface and total Itgb1 levels in USP12/46/SNX17-triple (t)KO fibroblast clones were further decreased compared to those in the USP12/46-dKO fibroblasts, suggesting that the DUB complex and SNX17 act independently of each other in maintaining Itgb1 levels (Fig. 3A-C). Re-expression of USP12 in USP12/46-dKO or SNX17 in SNX17-KO fully restored the surface levels of Itgb1 (Fig. 3D). Co-expression of USP12 and SNX17 fully restored the Itgb1 levels in tKO fibroblasts, whereas separate expression of USP12 or SNX17 in tKO fibroblasts failed to normalize Itgb1 surface levels (Fig. 3D).

**Figure 3.**
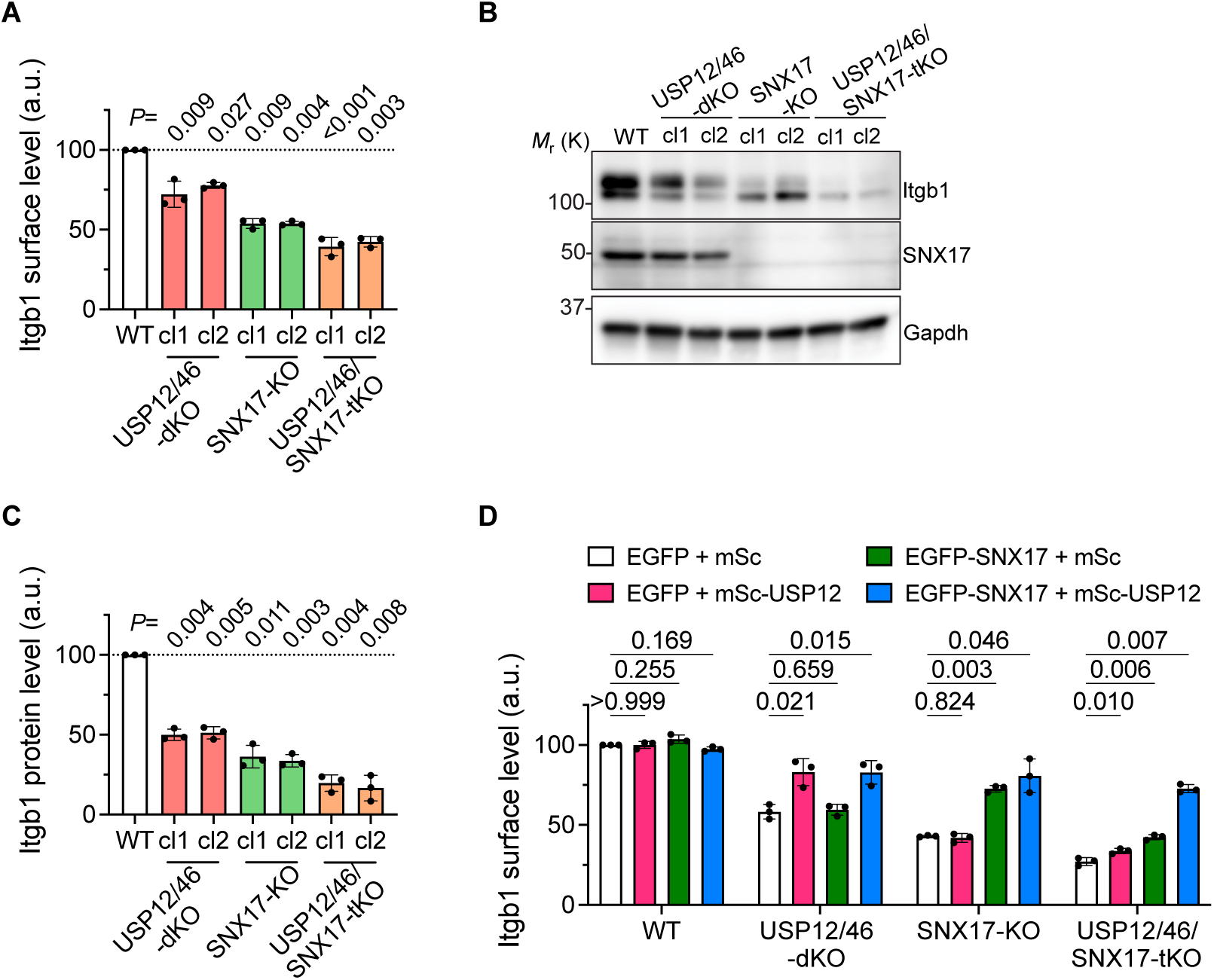
The USP12/46-WDRs complex stabilizes Itgb1 protein levels in a SNX17-independent manner. **(A-C)** Itgb1 surface levels determined by flow cytometry (**A**) and Itgb1 and SNX17 protein levels in cell lysates determined by WB (**B**) with densitometric quantification (**C**) in WT, USP12/46-dKO, SNX17-KO, and USP12/46/SNX17-tKO fibroblasts. Gapdh served as a loading control. Statistical analysis was carried out by RM one-way ANOVA with Dunnett’s multiple comparison test. Data are shown as Mean±SD, n=3 independent experiments. **(D)** Itgb1 surface levels in WT, USP12/46-dKO, SNX17-KO, and USP12/46/SNX17-tKO fibroblasts transiently expressing indicated combinations of constructs determined by flow cytometry. Statistical analysis was carried out by RM two-way ANOVA with Dunnett’s multiple comparison test. Data are shown as Mean±SD, n=3 independent experiments.

### The USP12/46-WDRs complex prevents lysosomal degradation of Itgb1

Next, we investigated the mechanism underlying the downregulation of Itgb1 expression in USP12/46-dKO cells. A role for *Itgb1* mRNA transcript stability in regulating Itgb1 protein levels could be excluded as no difference in *Itgb1* mRNA levels were found between USP12/46-dKO and WT fibroblasts (Fig. 4A). Cycloheximide (CHX) chase assays, which allow to compare the degradation kinetics of proteins, revealed accelerated Itgb1 protein degradation in USP12/46-dKO compared to WT fibroblasts (Fig. 4B, C). Furthermore, surface biotinylation followed by capture ELISA (Bottcher *et al*., 2012) showed a significantly reduced Itgb1 surface stability in USP12/46-dKO fibroblasts compared to WT fibroblasts. The Itgb1 protein half-life was approximately 12 hours in USP12/46-dKO fibroblasts and more than 20 hours in WT fibroblasts (Fig. 4D). The reduced Itgb1 surface stability in USP12/46-dKO fibroblasts was restored upon re-expression of USP12^WT^ but not upon re-expression of the catalytically inactive USP12^C48S^ (Fig. 4E). We also found that the internalization kinetics of Itgb1 was similar between USP12^WT^ and USP12^C48S^ re-expressing USP12/46-dKO fibroblasts (Fig. 4F), whereas the recycling rate of Itgb1 was reduced in USP12^C48S^ expressing fibroblasts compared to USP12^WT^ expressing fibroblasts (Fig. 4G). These data indicate that the catalytic activity of the USP12/46-WDRs complex stabilizes the surface as well as total Itgb1 protein levels by enabling the recycling of internalized Itgb1 to the cell surface.

**Figure 4.**
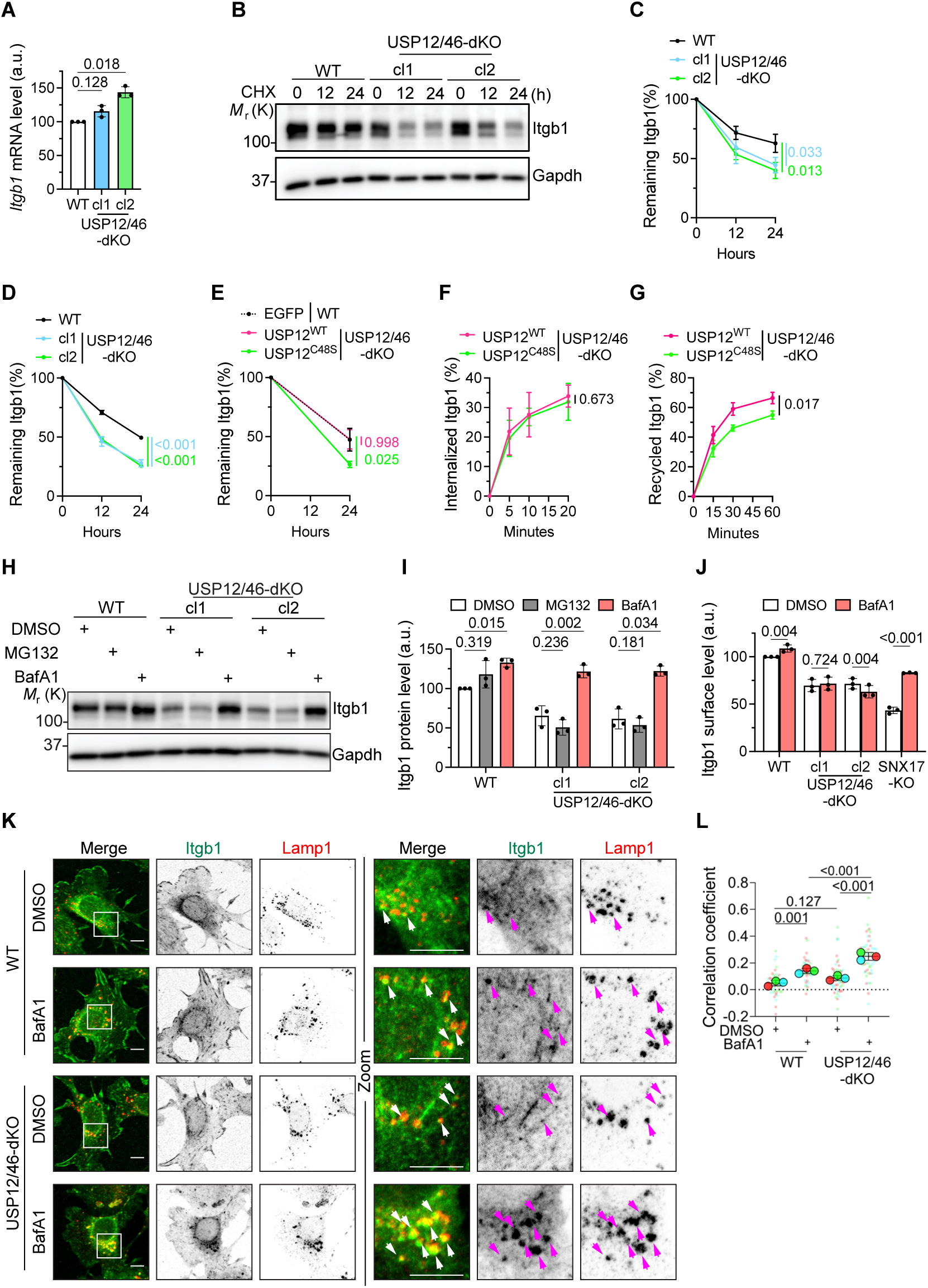
USP12/46-WDRs complex prevents lysosomal degradation of Itgb1. **(A)** *Itgb1* mRNA levels in WT and USP12/46-dKO fibroblasts determined by qPCR. Statistical analysis was carried out by RM one-way ANOVA with Dunnett’s multiple comparison test. Data are shown as Mean±SD, n=3 independent experiments. **(B, C)** WB (**B**) and densitometric quantification (**C**) of Itgb1 protein levels in WT and USP12/46-dKO fibroblasts at indicated time points after cycloheximide (CHX) treatment (5 μg/ml). Gapdh served as a loading control. Statistical analysis was carried out by ordinary one-way ANOVA with Dunnett’s multiple comparison test. Data are shown as Mean±SD, n=3 independent experiments. **(D)** Quantification of surface Itgb1 degradation kinetics in WT and USP12/46-dKO fibroblasts. The amount of Itgb1 remaining over indicated times were measured by capture-ELISA. Statistical analysis was carried out by ordinary one-way ANOVA with Dunnett’s multiple comparison test. Data are shown as Mean±SD, n=3 independent experiments. **(E)** Quantification of surface Itgb1 degradation kinetics in WT fibroblasts stably expressing EGFP, and USP12/46-dKO fibroblasts stably expressing EGFP-USP12^WT^ or EGFP-USP12^C48S^ determined by capture-ELISA. Statistical analysis was carried out by ordinary one-way ANOVA with Dunnett’s multiple comparison test. Data are shown as Mean±SD, n=3 independent experiments. **(F, G)** The internalization rate (**F**) and the recycling rate (**G)** of Itgb1 in USP12/46-dKO fibroblasts stably expressing EGFP-USP12^WT^ or EGFP-USP12^C48S^ determined by capture-ELISA. Statistical analysis was carried out by two-sided Welch’s *t*-test. Data are shown as Mean±SD, n=3 independent experiments. **(H, I)** WB (**H**) with densitometric quantification (**I**) of Itgb1 protein levels in lysates of WT and USP12/46-dKO fibroblasts treated with DMSO, MG132 (0.5 uM) or BafA1 (40 nM) for 9 hours. Gapdh served as a loading control. Statistical analysis was carried out by RM two-way ANOVA with Dunnett’s multiple comparison test. Data are shown as Mean±SD, n=3 independent experiments. **(J)** Itgb1 surface levels in WT, USP12/46-dKO and SNX17-KO fibroblasts treated with DMSO or BafA1 (40 nM) for 9 hours determined by flow cytometry. Statistical analysis was carried out by RM two-way ANOVA with Šidák’s multiple comparison test. Data are shown as Mean±SD, n=3 independent experiments. **(K)** Representative immunofluorescence (IF) images of Itgb1 and Lamp1 in WT and USP12/46-dKO fibroblasts treated with DMSO or BafA1 (100 nM) for 3 hours. Arrowheads show the accumulation of Itgb1 in Lamp1-positive endo/lysosomes. Boxes indicate magnified cell regions displayed in the Zoom panel. Sum intensity projections from confocal stacks are presented. Scale bar, 10 µm. **(L)** Superplots showing the Pearson correlation coefficients (PCC) between Itgb1 and Lamp1 in WT and USP12/46-dKO fibroblasts. Statistical analysis was carried out by RM two-way ANOVA with Šidák’s multiple comparison test. Data are shown as Mean±SD, n=3 independent experiments; 49 cells were analyzed in DMSO-treated WT cells, 43 in BafA1-treated WT cells, 52 in DMSO-treated USP12/46-dKO cells and 45 in BafA1-treated USP12/46-dKO cells.

To determine the pathway through which Itgb1 is degraded in the absence of USP12/46, we treated USP12/46-dKO fibroblasts with MG132 or Bafilomycin A1 (BafA1). Whereas inhibition of the proteasome with MG132 did not restore Itgb1 levels, inhibition of the lysosome with BafA1 restored total Itgb1 levels in cell lysates (Fig. 4H, I). Surprisingly, however, BafA1 treatment did not restore the surface levels of Itgb1 in USP12/46-dKO fibroblasts (Fig. 4J). To determine the subcellular localization of Itgb1 in WT and USP12/46-dKO fibroblasts, we immuno-stained fixed cells and found Itgb1 in FAs, ER, and a few intracellular puncta in both, WT and USP12/46-dKO fibroblasts. Following BafA1 treatment, we observed large Itgb1-positive puncta co-stained with the late endosome/lysosome marker Lamp1 in USP12/46-dKO fibroblasts, which were rarely observed in WT fibroblasts (Fig. 4K). An increased Pearson correlation coefficient (PCC) of Itgb1 with Lamp1 in USP12/46-dKO fibroblasts confirmed the increased endo/lysosomal localization of Itgb1 in USP12/46-dKO compared to WT fibroblasts (Fig. 4L). Collectively, these findings indicate that USP12/46 depletion leads to an increased lysosomal targeting of Itgb1, resulting in degradation, impaired recycling and reduced Itgb1 surface levels.

### The USP12/46-WDRs complex prevents ESCRT-mediated degradation of Itgb1

Since BafA1 prevents the degradation of proteins in lysosomes by blocking the vacuolar-type ATPase and the lysosomal acidification (Wang *et al*, 2021), we hypothesized that the absent Itgb1 deubiquitination in USP12/46-dKO cells promotes ESCRT binding, internalization of Itgb1 via intraluminal vesicles (ILVs) and Itgb1 degradation. To test this hypothesis, we conjointly depleted ESCRT-0 component HGS and ESCRT-I component TSG101 by siRNAs (ESCRT-KD) in WT and USP12/46-dKO fibroblasts and found that Itgb1 levels and stability increased in both, WT^ESCRT-KD^ and USP12/46-dKO^ESCRT-KD^ fibroblasts (Fig. 5A and Fig. EV5A-C). Immunoprecipitation of endogenous Itgb1 followed by WB for ubiquitinated Itgb1 with anti-Ub antibody revealed elevated poly-ubiquitination of Itgb1 in USP12/46-dKO^ESCRT-KD^ fibroblasts compared to WT^ESCRT-KD^ fibroblasts (see smears between 130-200 kDa in Fig. 5B; Fig. EV5D). The increased Itgb1 ubiquitination in USP12/46-dKO^ESCRT-KD^ fibroblasts was confirmed by capturing ubiquitinated proteins using ubiquitin-trap beads and subsequently probing the gel separated proteins with a polyclonal anti-Itgb1 antibody (Fig. 5C and Fig. EV5E).

**Figure 5.**
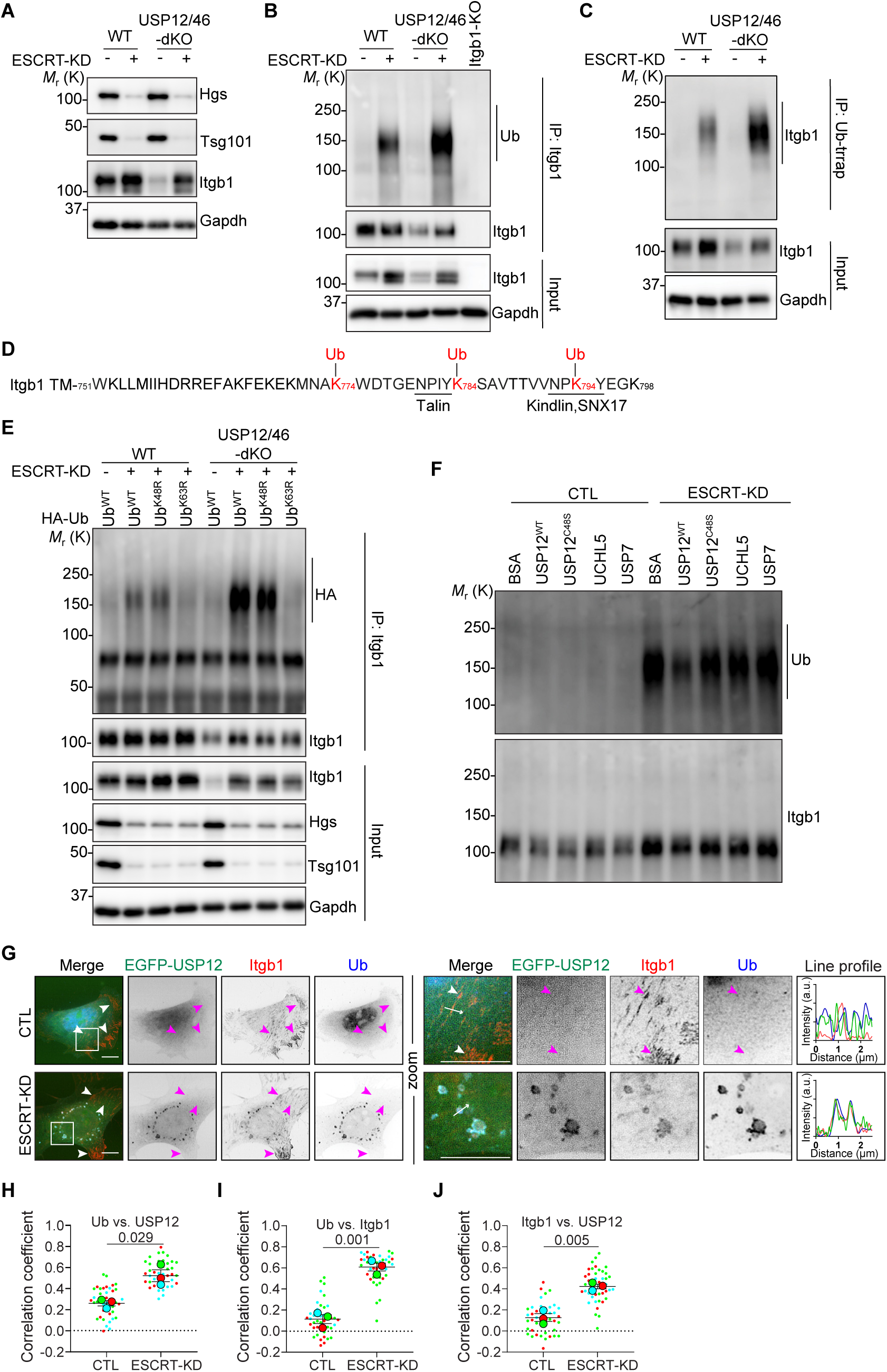
The USP12/46-WDRs complex deubiquitinates Itgb1 and prevents Itgb1 degradation via the ESCRT pathway. **(A)** WB of Itgb1 protein levels in WT and USP12/46-dKO fibroblasts transfected with control non-targeting siRNA (CTL) or siRNAs targeting jointly HGS and TSG101 (ESCRT-KD). Gapdh served as a loading control. Representative images from 3 independent experiments are shown. **(B)** IP of denatured Itgb1 from WT, USP12/46-dKO and Itgb1-KO fibroblast lysates with or without ESCRT-KD and analyzed by WB for Ubiquitin (Ub) and Itgb1. Itgb1-KO served as a negative control. Gapdh served as a loading control. Representative images from 3 independent experiments are shown. **(C)** IP of ubiquitinated proteins in WT and USP12/46-dKO fibroblasts with or without ESCRT-KD and analyzed by WB for Itgb1. Gapdh served as a loading control. Representative images from 3 independent experiments are shown. **(D)** Amino acid sequence of Itgb1 cytoplasmic tail. The ubiquitin-conjugated lysines detected by MS are colored in red. The underlined regions indicate the NPxY motifs that bind to Talin, Kindlin and SNX17. See also supplementary table S1. **(E)** IP of denatured Itgb1 from WT and USP12/46-dKO fibroblasts overexpressing HA-tagged Ub^WT^, Ub^K48R^ or Ub^K63R^ treated with or without ESCRT-KD siRNAs and analyzed by WB for HA and Itgb1. Gapdh served as a loading control. Representative images from 3 independent experiments were shown. **(F)** *In vitro* deubiquitination assay using recombinant WDR48-WDR20-USP12 (WT or C48S) complex, UCHL5 or USP7 and ubiquitinated Itgb1 enriched from USP12/46-dKO^ESCRT-KD^ fibroblast cell lysate followed by WB for ubiquitin. BSA served as a negative control. Representative images from 3 independent experiments were shown. **(G)** Representative Structured Illumination Microscopy (SIM) images of Itgb1 and Ub in USP12/46-dKO fibroblasts stably expressing EGFP-USP12 treated with or without ESCRT-KD siRNAs. Boxes indicate magnified cell regions displayed in the Zoom panel. Arrowheads show the Itgb1-labeled focal adhesion sites and arrows show the direction of line profiles. Scale bar, 10 µm. **(H-J)** Superplots showing the PCC between Ub and EGFP-USP12 (**H**), Ub and Itgb1 (**I**), and Itgb1 and EGFP-USP12 (**J**) in USP12/46-dKO fibroblasts stably expressing EGFP-USP12 and treated with or without ESCRT-KD siRNAs. Statistical analysis was carried out by two-sided Welch’s *t*-test. Data are shown as Mean±SD, n=3 independent experiments, in total 42 cells per condition.

To identify the lysine residues in the Itgb1 tail that are ubiquitinated, we immunoprecipitated Itgb1, treated the precipitate with trypsin and used MS to identify peptides containing a Gly-Gly motif, which is a remaining signature of an Ub-conjugated site (Xu *et al*, 2010). The experiment identified lysine-774 (K774), K784 and K794 modified by ubiquitin (Fig. 5D, Supplementary table S1). Interestingly, K794 is located within the Kindlin- and SNX17-binding NPK_794_Y motif and K784 is adjacent to the Talin-binding NPIY_783_ motif.

The ubiquitin (Ub) chain linkage specificity determines fate and function of polyubiquitinated proteins (Miranda & Sorkin, 2007; Swatek & Komander, 2016). Lysine-63 (K63)-linked polyUb chains are preferentially associated with ESCRT-mediated lysosomal degradation, while lysine-48 (K48)-linked chains lead to proteasomal degradation (Nathan *et al*, 2013; Strickland *et al*, 2022). To inhibit K48- or K63-mediated Ub conjugation we overexpressed ubiquitin mutants carrying K48R or K63R substitutions (Lim *et al*, 2005) and concomitantly depleted WT and USP12/46-dKO fibroblasts with ESCRT-KD siRNAs to enrich for ubiquitinated Itgb1 (Fig. 5E and Fig. EV5F). The experiment revealed an increase of ubiquitinated Itgb1 in cells expressing Ub^WT^ or Ub^K48R^, but not Ub^K63R^. The levels of Itgb1 ubiquitinated with Ub^WT^ and Ub^K48R^ were higher in USP12/46-dKO^ESCRT-KD^ compared to WT^ESCRT-KD^ fibroblasts, which indicates that K63-mediated polyUb chain modification destines Itgb1 for lysosomal degradation.

To further confirm the role of the USP12/46-WDRs complex in the deubiquitination of Itgb1, we performed an *in vitro* deubiquitination assay with recombinantly produced ternary wildtype USP12^WT^-WDR48-WDR20 or catalytically inactive USP12^C48S^-WDR48-WDR20 complexes (Fig. 5F). To enrich for ubiquitinated Itgb1, fibroblasts were first depleted with ESCRT-KD siRNAs and then Itgb1 was immunoprecipitated from cell lysates and incubated with the recombinant USP12^WT^-WDRs or the recombinant USP12^C48S^-WDRs complex (Fig. EV5G). Whereas USP12^WT^-WDRs reduced Itgb1 ubiquitination, the catalytically inactive USP12^C48S^-WDRs did not. Furthermore, neither recombinant UCHL5 nor USP7, which are integrin-unrelated DUBs and were used as controls, were able to reduce Itgb1 ubiquitination (Fig. 5F and Fig. EV5H). We also found that the EGFP-tagged USP12 expressed in USP12/46-dKO^ESCRT-KD^ fibroblasts colocalized with Itgb1 and ubiquitin on the limiting membrane of enlarged endosomes and was absent from Itgb1-positive focal adhesions (FAs) (Fig. 5G-J). Enlarged endosomes were not observed in control siRNA transfected USP12/46-dKO fibroblasts. Altogether, these data indicates that the USP12/46-WDRs complex colocalizes with Itgb1 on endosomes and deubiquitinates Itgb1, which in turn prevents ESCRT-mediated Itgb1 degradation.

Itgb1 crosslinking followed by IP and MS excluded a direct association between Itgb1 and the USP12/46-WDRs complex (Supplementary table S2). Since EGFP-USP12 was absent from FAs (Fig. 5G) and the members of the USP12/46-WDRs complex were also undetectable in the integrin adhesome (Horton *et al*, 2015; Kuo *et al*, 2011; Schiller *et al*, 2011), we conclude that the USP12/46-WDRs complex binds Itgb1 on endosomes and not in FAs.

If USP12/46-WDRs complex-mediated deubiquitination prevents Itgb1 degradation, non-ubiquitinable α5β1 integrins in which the lysine residues in the cytoplasmic tails of the α5 and β1 subunits were replaced with arginine residues should also escape ESCRT-mediated degradation and be recycled to the cell surface. To test this hypothesis, we generated Itgb1-KO fibroblasts expressing endogenous USP12/46 (Itgb1-KO^WT^) or lacking USP12/46 (Itgb1-KO^USP12/46-dKO^) and expressed α5^WT^β1^WT^ or α5^KR^β1^KR^, in which the 4 lysine residues in the Itga5 tail and the 8 lysine residues in the Itgb1 tail were substituted for arginine residues (Bottcher *et al*., 2012) (Fig. 6A). BafA1 treatment to block lysosomal degradation followed by immunostaining revealed that α5^WT^β1^WT^ accumulated in Lamp1^+^ endo/lysosomes in Itgb1-KO^USP12/46-dKO^ fibroblasts, whereas α5^KR^β1^KR^ did not accumulate (Fig. 6B). PCC confirmed the lower colocalization between Lamp1 and α5^KR^β1^KR^ compared to Lamp1 and α5^WT^β1^WT^ (Fig. 6C). Consistent with this observation, BafA1 treatment increased the surface levels of α5^KR^β1^KR^ and to a much lower extent of α5^WT^β1^WT^ in Itgb1-KO^USP12/46-dKO^ fibroblasts (Fig. 6D). Moreover, treatment with ESCRT-KD siRNAs significantly increased the surface levels of α5^WT^β1^WT^ but barely of α5^KR^β1^KR^ in Itgb1-KO^USP12/46-dKO^ fibroblasts, indicating that the ESCRT binds Ub^K63^ modified Itgb1 tails, which in turn leads to Itgb1 degradation and thereby prevents Itgb1 from being recycled to cell surface (Fig. 6E).

**Figure 6.**
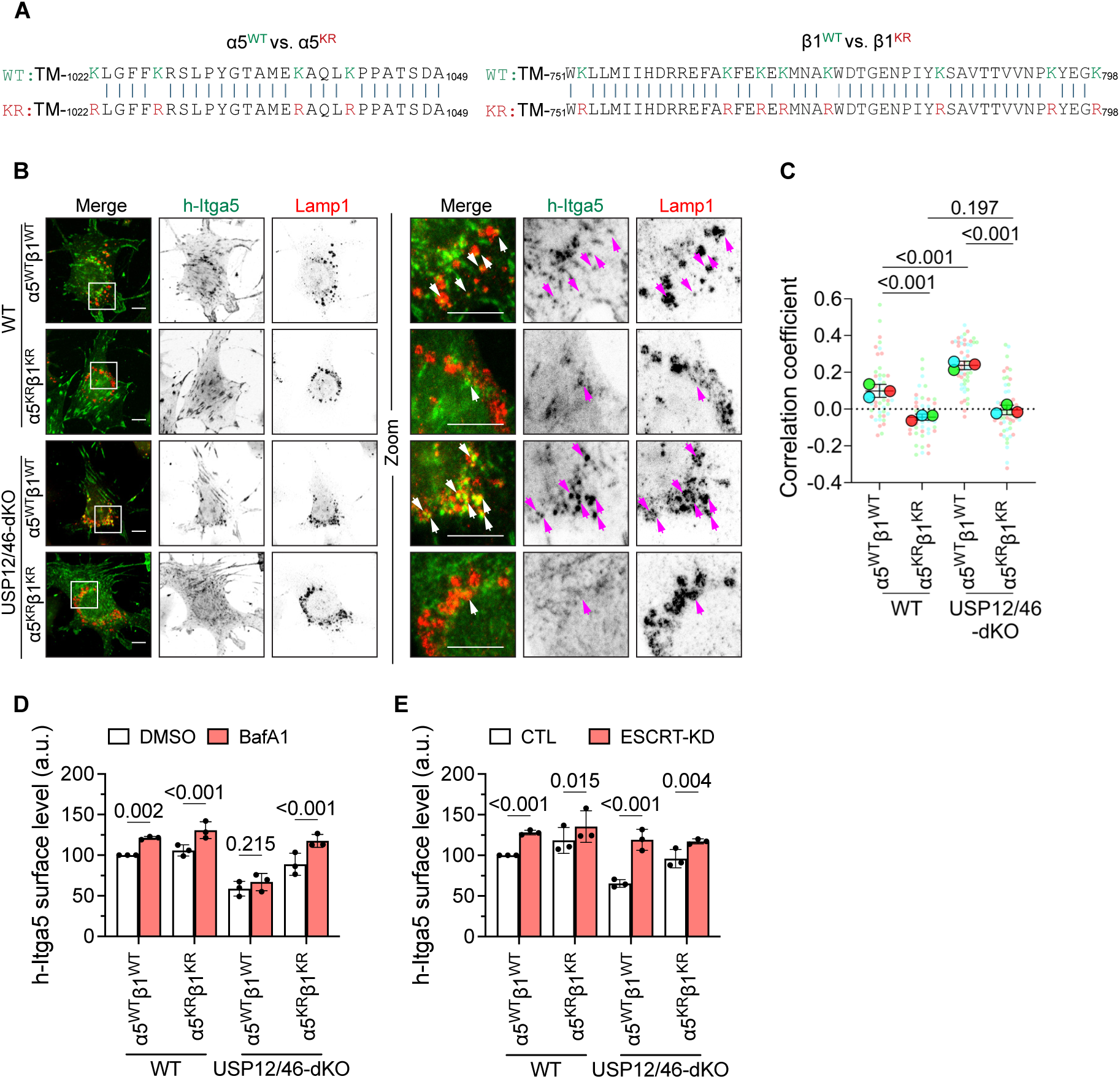
Itgb1 ubiquitination is required for Itgb1 sorting into the endosomal lumen. **(A)** Amino acid sequence alignments of the cytosolic tails of WT α5β1 integrin (α5^WT^β1^WT^) and mutant α5^KR^β1^KR^ where lysine residues were substituted for non-ubiquitinable arginine residues. **(B)** Representative IF images of exogenous human Itga5 (h-Itga5) and Lamp1 in BafA1-treated Itgb1-KO^WT^ and Itgb1-KO^USP12/46-dKO^ fibroblasts. Itgb1-KO cells were retrovirally transduced with human α5^WT^β1^WT^ or α5^KR^β1^KR^ integrins. A human specific Itga5 antibody was used for IF. Boxes indicate magnified cell regions displayed in the Zoom panel. Arrowheads indicates the colocalization of h-Itga5 with Lamp1. Sum intensity projections from confocal stacks are shown. Scale bar, 10 µm. **(C)** Superplots showing the PCC between Itgb1 and Lamp1 in fibroblasts as described above. Statistical analysis was carried out by ordinary two-way ANOVA with Šidák’s multiple comparison test. Data are shown as Mean±SD, n=3 independent experiments; 48 cells were analyzed in α5^WT^β1^WT^-expressing Itgb1-KO^WT^ cells, 44 in α5^KR^β1^KR^-expressing Itgb1-KO^WT^ cells, 52 in α5^WT^β1^WT^-expressing Itgb1-KO^USP12/46-dKO^ cells and 55 in α5^KR^β1^KR^-expressing Itgb1-KO^USP12/46-dKO^ cells . **(D)** Human Itga5 surface levels in DMSO- or BafA1-treated cells as described above. Statistical analysis was carried out by RM two-way ANOVA with Šidák’s multiple comparison test. Data are shown as Mean±SD, n=3 independent experiments. **(E)** Human Itga5 surface levels in cells as described above treated with or without ESCRT-KD siRNAs. Statistical analysis was carried out by RM two-way ANOVA with Šidák’s multiple comparison test. Data are shown as Mean±SD, n=3 independent experiments.

### The stabilization of Itgb1 by USP12/46-WDRs promotes integrin functions

In line with the increased ubiquitination, elevated degradation and decreased levels of cell surface Itgb1 associated with loss of USP12/46 expression, we observed an impaired adhesion, spreading and *in vitro* wound healing-based migration of USP12/46-dKO fibroblasts and transwell migration of USP12/46-dKO MDA-MB-231 cells (Fig. 7A-E and Fig. EV6A). Furthermore, USP12/46-dKO MDA-MB-231 cells showed reduced invasion through Matrigel-coated transwell filters compared to WT MDA-MB-231 cells (Fig. 7F, G). In line with these findings, analysis of the TCGC BRCA database linked high expression of USP12 and USP46 with reduced overall survival and progression free intervals of breast cancer patients (Fig. 7H, I and Fig. EV6B-E). These findings indicate that the Itgb1 deubiquitinating and stabilizing function of the USP12/46-WDRs complex is essential for basic integrin functions such as adhesion, spreading and migration. Furthermore, if their levels surpass a certain threshold such as in cancer Itgb1 levels rise, which is advantageous for invading cancer cells.

**Figure 7.**
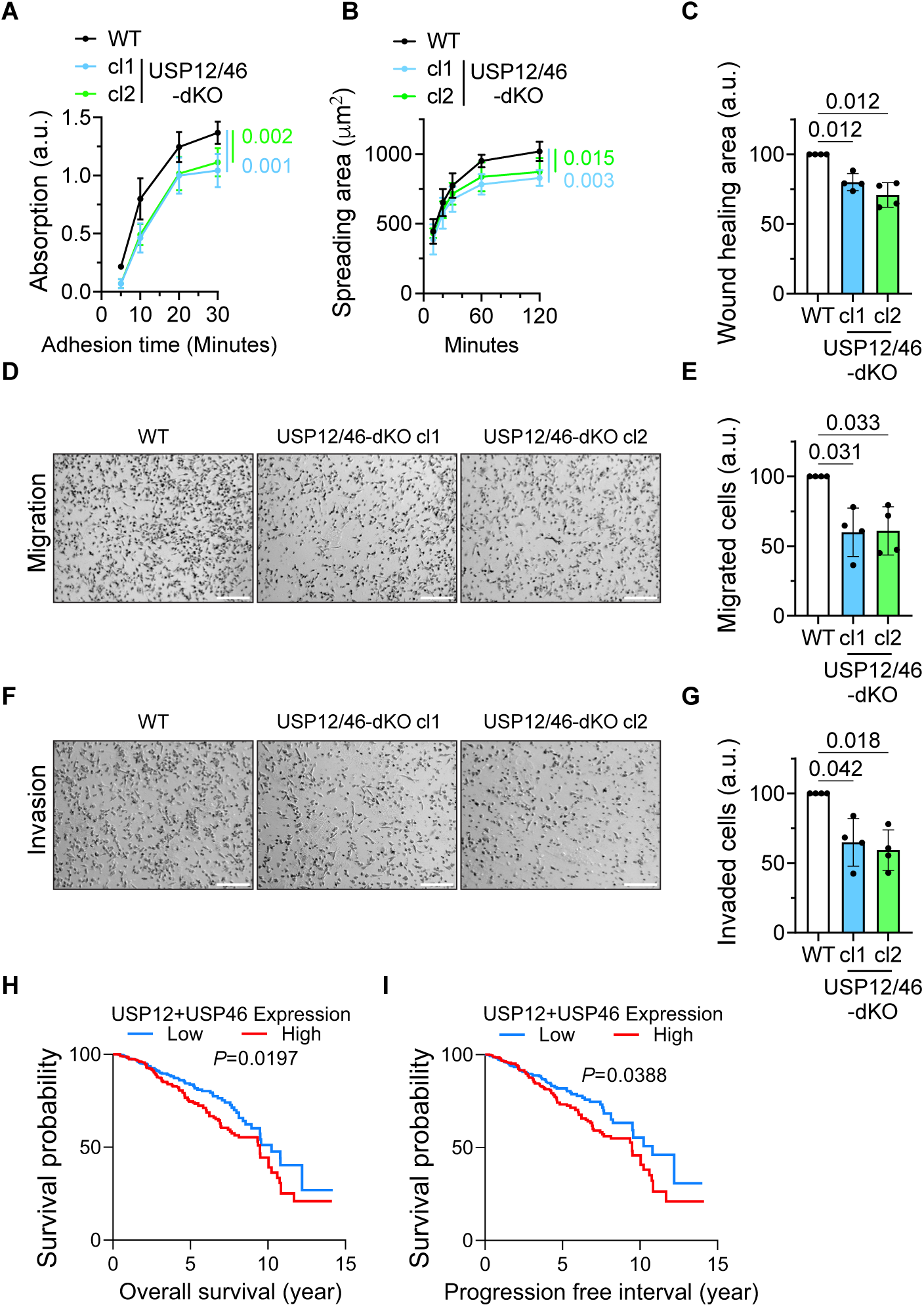
The USP12/46-WDRs complex promotes cancer cell migration and invasion. **(A)** Quantification of cell adhesion of WT and USP12/46-dKO fibroblasts at indicated time points after seeding on FN-coated plates. Statistical analysis was carried out by RM one-way ANOVA with Dunnett’s multiple comparison test at the 30-minute time point. Data are shown as Mean±SD, n=4 independent experiments. **(B)** Cell spreading area of WT and USP12/46-dKO fibroblasts on FN-coated glass surface shown at indicated time points after seeding. Statistical analysis was carried out by RM one-way ANOVA with Dunnett’s multiple comparison test at the 120-minute time point. Data are shown as Mean±SD, n=4 independent experiments. **(C)** Normalized wound healing area of WT and USP12/46-dKO fibroblasts on FN-coated plates after 12 hours. Statistical analysis was carried out by RM one-way ANOVA with Dunnett’s multiple comparison test. Data are shown as Mean±SD, n=4 independent experiments. **(D, E)** Representative images (**D**) and quantification (**E**) of WT and USP12/46-dKO MDA-MB-231 cells upon migration through transwell inserts 16 hours after seeding. Scale bar, 200 µm. Statistical analysis was carried out by RM one-way ANOVA with Dunnett’s multiple comparison test. Data are shown as Mean±SD, n=4 independent experiments. **(F, G)** Representative images (**F**) and quantification (**G**) of WT and USP12/46-dKO MDA-MB-231 cells upon migration through Matrigel-coated transwell inserts 16 hours after seeding. Scale bar, 200 µm. Statistical analysis was carried out by RM one-way ANOVA with Dunnett’s multiple comparison test. Data are shown as Mean±SD, n=4 independent experiments. **(H, I)** Kaplan Meier-plot of overall survival (**H**) **a**nd progression free interval (**I**) of breast cancer patients with high (red line) or low (blue line) combined total *USP12* and *USP46* gene expression levels. The GDC TCGA dataset obtained from the UCSC Xena project (Goldman *et al*, 2020) was used. Two-group risk model with cut-off at the median was applied. *P*-values were calculated by Log-rank test.

## Discussion

The steady-state surface level of Itgb1 underlies a tightly regulated decision-making process, which routes internalized Itgb1 either to lysosomes for degradation or to recycling endosomes for their reuse on the cell surface. The decision depends on ubiquitin moieties that are attached to the Itgb1-tail upon internalization and are either retained and recognized by the ESCRT machinery or removed by DUBs. USP9X, a DUB best known for its role in mitosis and DNA repair (Meng *et al*, 2023), was shown to remove ubiquitin from α5-integrin tails and stabilize α5β1 integrins in starved cells treated with soluble FN (Kharitidi et al., 2015). Since we excluded an involvement of USP9X in controling α5β1 integrin in cells cultured under steady state conditions, we decided to perform a Crispr/Cas9-based genetic screen and a BioID-based proximitome screen to identify DUB(s) that stabilize the steady-state surface levels of Itgb1.

Our screens identified the ternary DUB complex consisting of the deubiquitinase USP12 (or its paralog USP46) and two accessory proteins, WDR48 and WDR20, which form a complex that facilitates the stabilization of Itgb1 in sub-confluent HAP1 cells, MDA-MB-231 cells and mouse fibroblasts. In search for a mechanistic explanation for the Itgb1 protein stabilization, we found that the activity of the USP12/46-WDRs complex removes ubiquitin from Itgb1 at early endosomal membranes and thereby, inhibits ESCRT-mediated sorting of Itgb1 into intraluminal vesicles (ILVs) and lysosomal degradation, and instead retrieves Itgb1 into recycling endosomes. Consistently, loss of the two DUBs, USP12 and USP46, which compensate each other, or of either WDR20 or WDR48, which facilitates the DUB activity, decreases the half-life of the Itgb1 protein from 24 to 12 hours. Importantly, our cell surface proteome analysis also revealed that the USP12/46-WDRs complex regulates not only the stabilization of Itgb1 but also of other surface proteins, including signaling receptors and solute transporters. This diverse group of surface proteins with very different cytoplasmic domains did not reveal a common domain or motif that may serve as direct or indirect binding site of the USP12/46-WDRs complex (Yu et al., unpublished), nor did our attempts to compare the sequences of the Itgb1 tail with those of the known nuclear and cytosolic substrates of USP12/46-WDRs (Niu *et al*, 2023). Immunostaining and proteomics analysis indicate that the USP12/46-WDRs complex-mediated Itgb1 deubiquitination occurs at endosomes, as neither our immunostainings nor previous studies on the adhesome composition found the complex in integrin adhesion sites.

Our findings also show that the three lysine residues, K774, K784 and K794 in the Itgb1-tail are ubiquitin-modified in the absence of USP12/46. The K784 is located adjacent to the membrane proximal NPIY motif that serves as binding site for Talins, and the K794 is located in the distal NPKY motif that serves as binding site for Kindlin and SNX17. It is conceivable that the ubiquitination of K784 and K794 couples integrin inactivation by compromising Talin and Kindlin binding and blockade of endosomal retrieval and recycling by compromising SNX17 binding with ESCRT-mediated degradation. Hence, Itgb1-tail ubiquitination may have a dual function by coordinating activity and stability of integrins.

USP12 and USP46 can functionally compensate for each other *in vitro*. The two DUBs are ubiquitously expressed, however, display distinct expression levels in different tissues. USP12 is prominently expressed in bone marrow, whereas USP46 in muscle and brain tissues (proteinatlas.org) (Uhlen *et al*, 2015). Despite the different abundance of USP46 and USP12 in tissues, USP12-as well as USP46-KO mice are viable and lack obvious developmental and postnatal defects (mousephenotype.org) (Groza *et al*, 2023). However, increased levels of USP12 and USP46 have been associated with the progressions of several cancers, including breast cancer, liver cancer and multiple myeloma (Niu *et al*., 2023). The diminished Itgb1 levels in USP12/46-dKO MDA-MB-231 cells severely impaired migration and invasion. Hence, the link between USP12/46 and Itgb1 stability may well have a contributory role for the course of different cancer entities, originating of both epithelial and blood origin, and call for the exploration of compounds that inhibit the activity of the DUB complex.

## Methods

All methods can be found in the accompanying Supplementary Methods file.

## Acknowledgment

We thank the sequencing, mass spectrometry and imaging facilities of the Max Planck Institute of Biochemistry for the invaluable support, and Dr. Arnoud Sonnenberg for critically reading the manuscript and discussions. This work was supported by the German Research Foundation (project numbers 452409123 and 537477296) to F.B. and the European Research Council (ERC) under the European Union’s Horizon 2020 research and innovation program (grant agreement No. 810104 – Point) and the Max Planck Society to R.F.

## Author contribution

Conceptualization: GW and RF; Investigation: KY, SG and GW; Resource: FB; Supervision: GW and RF; Funding Acquisition: RF; Writing: KY, GW and RF.

## Materials & Methods

### Cell culture

HAP1 cells (C859, Horizon Discovery) were cultured in IMDM medium (#31980030, Gibco) supplemented with 10% FBS (#A5256701, Gibco). RPE1 cells (CRL-4000, ATCC), Hela cells (CCL-2, ATCC), MDA-MB-231 cells (HTB-26, ATCC) and mouse kidney fibroblasts (Böttcher *et al*, 2012) in DMEM (#61965059, Gibco) supplemented with 10% FBS (#A5256701, Gibco). All cells were cultured at 37 °C with 5% CO_2_ and routinely tested for mycoplasma.

### CRISPR screen

The CRISPR screen for human DUBs was carried out as previously described (Paulmann *et al*, 2022) with slight modifications. Briefly, 3x10^6^ HAP1 cells lentivirally transduced (lentiCas9-blast) to stably express the Cas9 protein (Sanjana *et al*, 2014) and cultured in the presence of 8 ug/ml protamine sulfate were retrovirally transduced with pooled sgRNA library targeting 98 DUBs and 5 genes essential for cell survival with 3 sgRNAs per gene and contained 12 non-targeting sgRNAs (Paulmann *et al*., 2022). Mutagenized HAP1 cells were cultured for one week, sorted for GFP expression by flow cytometry and expanded for another week. Subsequently, 40x10^6^ cells were harvested, resuspended in 0.4 ml FACS buffer (PBS containing 2% FBS and 2.5 mM EDTA), stained with PE-labeled anti-Itgb1antibody (#303004, Biolegend) for 45 minutes on ice, washed twice with ice-cold PBS, fixed with BD Cytofix^TM^ Fixation Buffer (#554655, BD) for 10 minutes on ice and then for 10 minutes at room temperature (RT). The fixed cells were stored in FACS buffer supplemented with 0.01% sodium azide at 4 °C in the dark before cells were sorted with a FACSAriaIII flow cell sorter. Prior to loading onto the sorter, cells were stained with Hoechst 33343 (#H1399, Thermofisher) for 30 minutes on ice and haploid cells were gated based on the Hoechst signal followed by forward and sideward gating, and finally gating 5% high and 5% low PE-positive Itgb1 expressing cell populations.

Genomic DNA from the 5% high and 5% low populations was extracted followed by the amplification of the sgRNA cassettes by a two-step PCR approach and by adding adapters and sample barcodes for deep sequencing. PCR products were sequenced on an Illumina NextSeq 500 instrument and the fastq output files were analyzed on the Galaxy platform (Galaxy, 2022) using the instances at usegalaxy.eu. Reads were mapped into the sgRNA library and the count table were analyzed with MAGeCK packages. Figure was generated using the R software with the packages tidyverse, ggplot and ggrepel.

### Generation of cell lines

The Crispr-Cas9 technique was used to generate knockout (KO) clones of USP12, USP46, WDR48, WDR20 and SNX17 following published protocols (Ran *et al*, 2013). The vector pSpCas9(BB)-2A-Puro (PX459) V2.0 (a gift from Feng Zhang, Addgene plasmid # 62988) was used to express the gRNA and Cas9. The gRNA sequences and primers for genomic DNA amplification and sequencing are listed in Table 1.

**Table 1.**
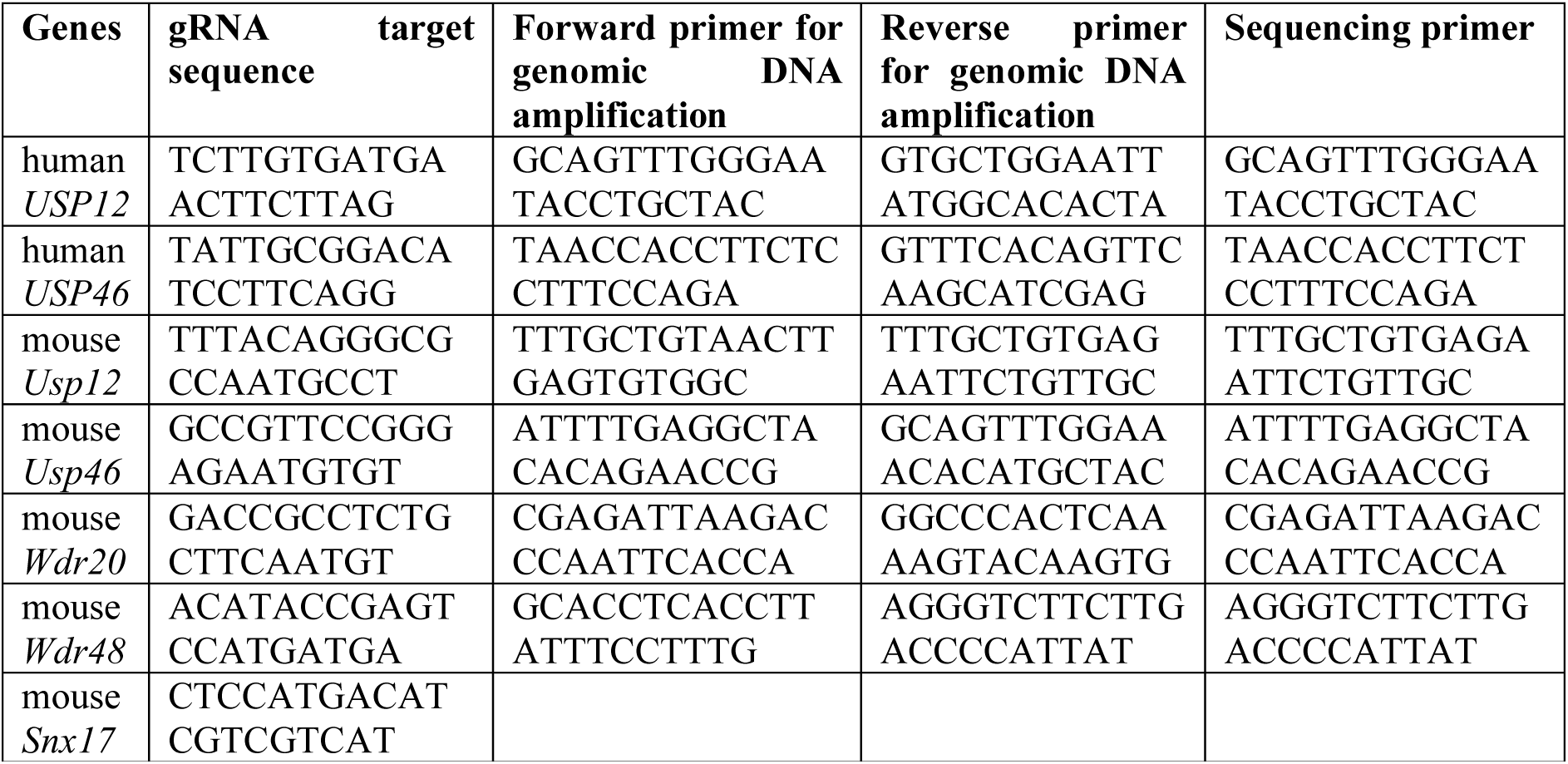
gRNA targeting sequences and primers for genomic PCR and Sanger sequencing for specified genes.

The gene disruptions were confirmed by sequencing the genomic DNA regions which were targeted by the gRNAs or Western blot if specific antibodies were available. Itgb1-KO fibroblasts were generated by adding adenoviral Cre to Itgb1-floxed fibroblasts derived from the kidneys of adult Itgb1-floxed mice (Böttcher *et al*., 2012).

To express ectopic integrin, Itgb1-null fibroblasts were retrovirally transduced with cDNAs encoding human Itgb1 and Itga5. WT or nullizygous USP12, USP46 and WDR48 cells were reconstituted by retrovirally transducing the cells with cDNA constructs tagged with EGFP, mScarlet or 3XFlag tags, respectively, as indicated in the Results section. To generate cells stably expressing miniTurbo-tagged Itga5, Itgb1-floxed fibroblasts were retrovirally transduced with the human Itga5 tagged with the miniTurbo DNA at the 3’-end.

### Plasmids and siRNAs

The mouse USP12, USP46, WDR48 cDNAs were reverse transcribed from mouse kidney fibroblasts and cloned into pEGFP-C1 and pRetroQ-C1 vectors (Clontech). The mouse WDR20 cDNA was prepared as described above and cloned into pEGFP-N1 (Clontech). The cDNAs encoding catalytically inactive USP12^C48S^ and USP46^C44S^ and binding deficient USP12^E190K^, USP12^V279A/F287A^, USP12^E190K/V279A/F287A^, WDR48^K214E/W256A/R272D^ and WDR20^F262A/W306A^ were generated by site-directed mutagenesis and cloned into pEGFP-C1 and pRetroQ-C1 vectors (Clontech). The cDNAs encoding WDR48^1-580^ and WDR48^1-359^ deletion mutants were amplified from the WDR48^WT^ cDNA and cloned into the pRetroQ-C1 vector. The Itgb1 and Itga5 cDNAs carrying lysine for arginine substitutions in the cytoplasmic domain (Itgb1^8xKR^ and Itga5^4xKR^) were previously generated by site-directed mutagenesis (Böttcher *et al*., 2012) and cloned into the HSC1 retroviral vector (a gift from James Ellis, Addgene plasmid # 58254). The cDNA encoding miniTurbo tagged Itga5 was generated by fusing the miniTurbo tag (a gift from Alice Ting, Addgene plasmid # 107168) in frame to the 3’ end of the human Itag5 cDNA (a gift from Rick Horwitz, Addgene plasmid # 15238) and subsequently cloned into pRetroQ-N1 vector. The cDNAs encoding Ub-WT (a gift from Ted Dawson, Addgene plasmid # 17608), Ub-K48R (a gift from Ted Dawson, Addgene plasmid # 17604) and Ub-K63R (a gift from Cecile Pickart, Addgene plasmid # 18898) were cloned into pEGFP-N1 vector (Clontech).

Depletion of USP9X was carried out with siRNAs pools targeting human (L-006099-00-0005, Dharmacon) and mouse USP9X (L-046869-01-0005, Dharmacon) in mouse fibroblasts, RPE-1, Hela and MDA-MB-231 cells. The transduced cells were detached, 48 hours later, washed twice with PBS, seeded on PLL-coated 12-well plates in DMEM supplemented with 10% FBS or in FN-coated 12-well plates in serum replacement medium (Benito-Jardon *et al*, 2017) containing 46.5% AIM-V medium (#12055, Gibco), 5% RPMI-1640 medium (#61870, Gibco), 1% Non-essential amino acid (NEAA,#11140, Gibco) and 47.5% DMEM medium (Nr.9007.1, Carl Roth) and cultured overnight at 37 °C. The surface levels of Itgb1 were assessed by flow cytometry and the total levels by Western blot. To disrupt the ESCRT function, a co-depletion was performed using siRNAs targeting HGS (L-055516-01-0005, Dharmacon) and TSG101 (L-049922-01-0005, Dharmacon) (Lobert *et al*, 2010). A non-targeting siRNA (D-001810-10-05, Dharmacon) was used as control.

### Transient transfection and viral transduction

Cells were transfected with plasmids using Lipofectamine 2000 or with siRNA using Lipofectamine RNAiMAX following the manufacturer’s protocol (Invitrogen). Viral transduction of expression constructs to generate stable cell lines was carried out as described previously (Theodosiou *et al*, 2016).

### Antibodies and reagents

The following antibodies were used for Western blot: α5 integrin (#4705, Cell Signaling Technology, 1:1,000), human β1 integrin (Clone 18, BD biosciences, 1:1,000), mouse β1 integrin (homemade, 1: 10,000) (Azimifar *et al*, 2012), GAPDH (CB1001, Calbiochem, 5,000), Flag (clone M2, Sigma, 1:1,000), GFP (A10262, Invitrogen, 1:1,000), SNX17 (10275-1-AP, Proteintech, 1:1,000), haemagglutin (HA)-tag (clone 3F10, Roche, 1:1,000), WDR48 (sc-514473, Santa Cruz Biotechnology, 1:500), WD20 (sc-100900, Santa Cruz Biotechnology, 1:500), HGS (10390-1-AP, Proteintech, 1:1,000), TSG101 (sc-7964, Santa Cruz Biotechnology, 1:1,000), ubiquitin (clone P4D1, Cell Signaling Technology, 1:1,000), RFP (PM005, MBL Life Science, 1:1,000).

The following antibodies were used for flow cytometry analysis: mouse β_1_ integrin (HMbeta1-1, Biolegend, 1:400), mouse α_5_ integrin (5H10-27, BD Pharmingen, 1:400), human β_1_ integrin (Ha2/5, BD Pharmingen, 1:400), human α_5_ integrin (IIA1, BD Pharmingen, 1:400).

The following antibodies were used for immunofluorescence (IF): mouse β_1_ integrin (homemade, 1:5,000) (Azimifar *et al*., 2012), human α_5_ integrin (VC5, BD biosciences, 1:400), mouse lamp1 (1D4B, DSHB, 1:400), ubiquitin (UBCJ2, Enzo Life Sciences, 1:200). Fluorophore-conjugated secondary antibodies: donkey anti-rat Alexa 488 (A21208, Invitrogen, 1:400), goat anti-rabbit Alexa 568 (A11036, Invitrogen, 1:400), donkey anti-mouse Alexa 568 (A10037, Invitrogen, 1:400), donkey anti-mouse Alexa 647 (A31571, Invitrogen, 1:400).

The following chemicals were used: TRITC-conjugated phalloidin (P1591, Sigma, 1:4,000 for staining); Hoechst 33342 (#H1399, ThermoFischer), N-ethylmaleimide (E-3876, Sigma); protease inhibitor cocktail (4693159001, Roche); DMSO (042780.AK, ThermoFisher); MG132 (474787, Sigma); bafilomycin A1 (BVT-0252, Adipogen); cycloheximide (sc-3508A, Santa Cruz); mytomycin C (M4287, Sigma); Poly-L-lysine solution (PLL, P4707, Sigma).

### Proximity-dependent biotin identification

To perform proximity-dependent biotin identification, Itgb1 floxed and Itgb1-KO cells stably expressing miniTurbo-tagged Itga5 were cultured in three 15-cm dishes, grown to 80% confluence, incubated with 50 µM biotin in PBS solution for 30 minutes at 37 °C, washed twice with PBS and then incubated with DMEM for a further 10 minutes. After cells were washed twice with PBS, lysates were generated with lysis buffer (0.1% SDS, 0.1% SDC, 1% Triton, 150 mM NaCl, 50 mM Tris, pH=8) containing protease inhibitor cocktail, then incubated with streptavidin magnetic beads (Cytiva) for 2.5 hours at 4 °C. The beads were washed three times with lysis buffer, once with PBS to remove detergent and then incubated in SDC buffer comprising of 1% sodium deoxycholate (SDC, Sigma), 40 mM 2-chloroacetamide (CAA, Sigma), 10 mM tris (2-carboxyethyl) phosphine (TCEP; ThermoFisher), and 100 mM Tris, pH 8.0 at 37 °C. After a 20-minute incubation at 37 °C, the samples were diluted at a 1:2 ratio with MS grade water (VWR), the proteins digested overnight at 37 °C with 0.5 µg of trypsin (Promega) and the resulting supernatant containing the peptide mixture harvested using a magnetic rack, acidified with trifluoroacetic acid (TFA; Merck) to achieve a final concentration of 1% and desalted using SCX StageTips. The samples were eluted, vacuum-dried and reconstituted in LC-MS grade water containing 0.1% formic acid before being loaded onto Evotips (Evotip Pure, Evosep).

A LC-MS/MS system coupled to a timsTOF Pro mass spectrometer (Bruker) was used to analyze the peptides. Evotips eluates were applied onto a 15-cm column (PepSep C18 15cmx15cm, 1.5 µm, Bruker Daltonics) utilizing the Evosep One HPLC system. The column temperature was set to 50 °C, peptide separation was achieved with the 30 SPD (samples per day) method, eluted and directly introduced into the timsTOF Pro mass spectrometer (Bruker Daltonics) via the nanoelectrospray interface. Data acquisition on the timsTOF Pro instrument was performed via the timsControl software. The MS functioned in data-dependent PASEF mode, performing one survey TIMS-MS scan and ten PASEF MS/MS scans per acquisition cycle. Analysis spanned a mass scan range of 100-1700 m/z and an ion mobility range from 1/K0 = 0.85 Vs cm-2 to 1.35 Vs cm-2, with uniform ion accumulation and ramp time of 100 milliseconds each in the dual TIMS analyzer, achieving a spectra rate of 9.43 Hz. Precursor ions suitable for MS/MS analysis were isolated within a 2 Th window for m/z < 700 and 3 Th for m/z > 700 by promptly adjusting the quadrupole position as precursors eluted from the TIMS device. Collision energy was adapted based on ion mobility, commencing at 45 eV for 1/K0 = 1.3 Vs cm-2 and decreasing to 27 eV for 0.85 Vs cm-2. Collision energies were interpolated linearly between these thresholds and remained constant above or below them. Uniquely charged precursor ions were filtered out employing a polygon filter mask, and supplementary m/z and ion mobility details were harnessed for ’dynamic exclusion’ to avert the re-evaluation of precursors once they attained a ’target value’ of 14500 a.u.

The MaxQuant computational platform (version 2.2.0.0) (Cox & Mann, 2008) was used to analyze the raw data with typical configurations tailored for Orbitrap or ion mobility data. Essentially, the peak list was cross-referenced against the Uniprot database of mus musculus (downloaded in 2023). Cysteine carbamidomethylation was designated as a fixed modification, while methionine oxidation and N-terminal acetylation were considered variable modifications. Protein quantification across runs was performed utilizing the MaxLFQ algorithm.

### Flow cytometry

To assess surface integrin levels, cells were trypsinized, washed with cold PBS, incubated with FACS antibodies diluted 1:400 in PBS supplemented with 1% BSA for 30 minutes on ice in the dark, washed with cold PBS and analyzed using the BD LSRFortessa^TM^ X-20 Cell Analyzer. The geometric mean fluorescence intensity of each sample was then evaluated using the FlowJo software version 10.10.

### Cell surface proteome

Cells were cultured in three independent 6-cm dishes at approximately 80% confluence. Cells were surface biotinylated with 0.2 mg/ml EZ-Link™ Sulfo-NHS-SS-Biotin (21217, ThermoFisher) in cold PBS for 30 minutes at 4 °C. Cell lysates were generated by incubating cells in lysis buffer (0.1% SDS, 0.1% SDC, 1% Triton, 150 mM NaCl, 50 mM Tris, pH=8) with protease inhibitor cocktail. The cell lysate was then incubated with Streptavidin Mag Sepharose beads (Cytiva#28985799) for 2.5 hours at 4 °C. The beads were washed three times with lysis buffer and once with PBS to remove detergent. Proteins on beads were digested by trypsin and the peptides were prepared as described above for proximity-dependent biotin identification. MDA-MB-231 samples were analyzed on the Brucker TimsTOF Pro with procedures and parameters as described above. Mouse fibroblast samples were analysed on an ThermoFisher QExactive HF mass spectrometer. Peptides were chromatographically separated using a 30-cm analytical column (inner diameter: 75 microns), packed in-house with ReproSil-Pur C18-AQ 1.9-micron beads from Dr. Maisch GmbH. This separation was achieved through a 60-minute gradient ranging from 8% to 30% buffer composed of 80% acetonitrile and 0.1% formic acid. The mass spectrometer operated in a data-dependent mode, with survey scans spanning from 300 to 1650 m/z at a resolution of 60,000 (at m/z = 200). It selectively picked up to 10 of the top precursors for fragmentation using higher energy collisional dissociation (HCD) with a normalized collision energy of 28. MS2 spectra were acquired at a resolution of 30,000 (at m/z = 200). Automatic gain control (AGC) targets for both MS and MS2 scans were set to 3E6 and 1E5, respectively, with maximum injection times of 100 ms for MS scans and 60 ms for MS2 scans. The raw data was analyzed as described above in proximity-dependent biotin identification.

### Quantitative PCR

Total RNA was extracted from fibroblasts using TRIzol™ Reagent (15596026, Invitrogen) following the manufacturer’s instructions. cDNA was synthesized using iScript™ cDNA Synthesis Kit (1708896, Bio-Rad), followed with real-time PCR using iQ SYBR Green Supermix (#1708880, Bio-Rad) on LightCycler® 480 (Roche). Primers for GAPDH: forward 5’-AGGTCGGTGTGAACGGATTTG-3’, reverse 5’-TGTAGACCATGTAGTTGAGGTCA-3’. Primers for Itgb1: forward 5’-ATGCCAAATCTTGCGGAGAAT-3’, reverse 5’-TTTGCTGCGATTGGTGACATT-3’.

### Western blot

Cells were lysed in RIPA buffer (0.1% SDS, 0.1% SDC, 1% Triton, 150 mM NaCl, 50 mM Tris, pH=8) containing protease inhibitor cocktail. Protein concentrations were determined using the BCA assay kit (#23225, ThermoFisher), lysates were boiled at 95 °C for 5 minutes in 1x Laemmli buffer, resolved on SDS-PAGE gels and transferred to 0.45 um PVDF membranes (IPVH00010, Millipore). After transfer, the membranes were briefly washed with PBS containing 0.1% Tween-20 (PBST) for 5 minutes before incubation with 5% BSA in PBST for 1 hour at RT, incubated with primary antibodies overnight at 4 °C , washed in PBST and then incubated with HRP-conjugated secondary antibodies for 1 hour at RT. A GE Amersham AI600 imager was used to detect the chemiluminescence generated upon addition of Immobilon Western Chemiluminescent HRP substrate (Millipore#WBKLS0500) to the membranes and Image Lab version 6.1 was used to quantify densitometries.

### Integrin degradation, internalization and recycling assays

These assays were carried out as previously described (Böttcher *et al*., 2012).

### Immunoprecipitation

The GFP-based immunoprecipitation (GFP IP) was carried out as previously described with slight modifications (Chen *et al*, 2022). In brief, cells were lysed in lysis buffer (50 mM Tris-HCl, 150 mM NaCl, 1 mM EDTA, 1% Triton, protease inhibitor cocktail) and the supernatant was incubated with GFP nano-trap beads (gta, Chromotek) for 2 hours at 4 °C. The beads were washed three times with the lysis buffer, then boiled in 2x Laemmli buffer, separated onto SDS–PAGE followed by WB for detecting the indicated proteins.

### Itgb1 *in vivo* crosslinking co-IP

Fibroblasts grown 15-cm dish to 80% confluence were siRNA-treated for 48 hours, washed with PBS twice, incubated on ice, then incubated with cross-linker (DSP, 0.1 mg/ml in PBS) solution or PBS for 30 minutes on ice, washed and quenched with quenching solution (50 mM Tris, pH 7.4, 150 mM NaCl, 1 mM MgCl_2_, 1 mM CaCl_2_). Cells were lysed in lysis buffer (50 mM Tris-HCl, 150 mM NaCl, 1 mM EDTA, and 1% Triton), the lysates were sonicated and cleared by centrifugation. The supernatant was incubated with anti-β1 integrin antibody (homemade) and protein A/G agarose beads (sc-2003, Santa Cruz) for 3 hours at 4 °C. The beads were washed with lysis buffer three times and one more time with PBS to remove the detergent. Peptides were prepared and processed on the MS as described above for proximity-dependent biotin identification. Raw data were analyzed using the Spectronaut 18.0 in directDIA+ (library-free) mode with the peak list was cross-referenced against the Uniprot database of mus musculus (downloaded in 2023). Cysteine carbamidomethylation was designated as a fixed modification, while methionine oxidation and N-terminal acetylation were considered variable modifications. Protein quantification across samples was achieved via label-free quantification (MaxLFQ) at the MS2 level.

### Ubiquitination assay

To measure endogenous ubiquitinated Itgb1, fibroblasts cultured in a 15-cm dish at approximately 80% confluence in the presence of 10% FBS were treated with non-targeting siRNA or siRNA simultaneously targeting mouse HGS and TSG101 (ESCRT-KD) for 48 hours. Cells were collected by scraping with PBS supplemented with 20 mM N-ethylmaleimide. Cell pellets were then lysed with 100 ul of 1% SDS in PBS and immediately boiled for 10 minutes to denature the proteins. The lysate was then diluted with 900 ul lysis buffer (1% Triton, 150 mM NaCl, 50 mM Tris, pH=8, 1 mM EDTA, 20 mM N-ethylmaleimide) supplemented with the protease inhibitor cocktail. The diluted lysate was further sonicated and cleared by centrifugation. The supernatant was then incubated with anti-β1 integrin antibody (homemade) and protein A/G agarose beads for 3 hours at 4 °C. The beads were then boiled in 2x Laemmli buffer, eluted and the elute was Western blotted for ubiquitin and Itgb1. Alternatively, cells were lysed in RIPA buffer (0.1% SDS, 0.1% SDC, 1% Triton, 150 mM NaCl, 50 mM Tris, 20 mM N-ethylmaleimide, pH=8) with protease inhibitor cocktail. The cell lysate was then incubated with the ubiquitin selector beads (N2510, NanoTag) for 3 hours at 4 °C to enrich for ubiquitinated proteins. Then the beads were boiled in 2x Laemmli buffer and the elute was subjected to SDS–PAGE followed by WB for Itgb1.

To assess the ubiquitination sites on Itgb1 cytoplasmic tail, the ubiquitin selector beads samples were prepared as above and peptides on the beads were prepared and analyzed by MS as described above for proximity-dependent biotin identification.

To determine the ubiquitin linkage specificity of Itgb1, fibroblasts were cultured in 15-cm dish to 80% confluence, transfected with constructs expressing HA tagged Ub^WT^, Ub ^K48R^ or Ub^K63R^ in the presence or absence of ESCRT-KD siRNA for 48 hours, harvested, followed by Itgb1 immunoprecipitation as described above. The beads were boiled in 2x Laemmli buffer, eluted and the elute was subjected to SDS–PAGE followed by WB for HA and Itgb1.

For the *in vitro* deubiquitination assay, USP12/46 dKO fibroblasts cultured in a 15-cm dish 80% confluence were treated with or without ESCRT-KD siRNA for 48 hours. Cells were harvested, Itgb1 was immunoprecipitated using the anti-β1 integrin antibody (homemade) coupled protein A/G agarose beads. The agarose beads were incubated with 100 nM recombinant USP12-WDRs complexes (generated as described below), recombinant UCHL5 (NBP1-72315, Novus Biologicals) or USP7 (E-519-025, Novus Biologicals) in a reaction buffer containing 40 mM Tris, 0.5 mM EDTA, 100 mM Nacl. 0.1% BSA, 1 mM TCEP (PH=7.4) for 30 minutes at 37 °C with constant shaking at 900 rpm in a thermal mixer. The beads were washed three times with washing buffer (1% Triton, 150 mM NaCl, 50 mM Tris, 1 mM EDTA, pH=8), boiled in 2x Laemmli buffer, eluted and the elute was subjected to SDS–PAGE followed by WB for ubiquitin and Itgb1.

### Expression and purification of recombinant proteins

Full-length recombinant WDR20 (1-569aa) N-terminally tagged with His6 was expressed in *E.coli.*, and WDR48 (1-580 aa) N-terminally tagged with Strep-tag II together with untagged USP12^WT^ and USP12^C48S^ (both 40-370aa) were expressed in insect cells as reported previously (Li *et al*, 2016) using the pCoofy expression vectors (a gift from Sabine Suppmann, Addgene plasmid # 43974) and purified to approximately 80% purity followed previously published protocols (Aretz *et al*, 2023). The purified USP12-WDR48 complex was incubated with WDR20 overnight to form the ternary DUB complex that was purified by size-exclusion chromatography. The purity of the recombinant proteins was verified by SDS-PAGE followed with Coomassie staining and MS.

### Immunofluorescence microscopy

Cells were grown overnight on FN-coated (5 µg/ml) coverslips that were kept in a 12-well plate (1x10^5^ per well), fixed with 4% PFA for 15 minutes at RT, and permeabilized with 0.1% Triton X-100 for 10 minutes at RT. Cells were blocked with 5% BSA for 1 hour at RT, incubated with primary antibodies overnight at 4 ℃, then with secondary antibodies (1:400 dilution) for 1 hour at RT and finally mounted in Elvanol No-Fade™ Mounting Medium.

Images were captured by the Zeiss LSM780 confocal laser scanning microscope or by the Zeiss Elyra PS.1 structured illumination microscope using ZEN software (Black version). Pearson correlation coefficient (PCC) analysis was performed using Fiji ImageJ software with the EzColocalization plug-in (Stauffer *et al*, 2018).

### Adhesion, spreading and wound healing assays

The adhesion and spreading assays were performed as previously described (Theodosiou *et al*., 2016) with slight modifications. Briefly, fibroblasts were starved for 4 hours in DMEM without FBS. Then the cells were trypsinized and incubated for a further 1 hour at 37 °C in DMEM supplemented with 3% BSA. The starved cells were then seeded on 96-well plates (40,000 per well) coated with either Poly-L-Lysine (PLL, P4707, sigma, 1:10 dilution) or fibronectin (5 µg/ml) or 3% BSA and incubated for 5, 10, 20 and 30 minutes at 37 °C, vigorously washed with PBS, fixed in 4% PFA, stained with 0.1% Crystal Violet solution and then dissolved in 2% SDS. The OD value of each well was acquired by a plate reader at 595 nm. Adhesion capacity was normalized using the equation: Normalized OD_595_= (OD_FN_-OD_BSA_)/ OD_PLL_.

For the spreading assay, the starved cells were seeded on FN-coated (5 µg/ml) glass coverslips and incubated at 37 °C for indicated time. Cells were then fixed in 4% PFA and stained with TRITC-conjugated phalloidin (1:4,000, 2 hours at RT) and Hoechst 33342 (1:10,000, 10 minutes at RT). Images were captured by the LSM780 confocal microscope and at least 45 cells per time point were analyzed for cell spreading area using Fiji ImageJ software.

For the wound healing assay, cells were seeded on FN-coated (5 µg/ml) 24-well plate containing a 2-well silicone insert (81176, ibidi) with a cell-free gap of approximately 500 µm overnight (1x10^4^ cells per well). Then the insert was removed, cells were briefly washed with PBS to remove cell debris and allowed to migrate for 12 hours at 37 °C with 5% CO_2_ in DMEM supplemented with mytomycin C (5 µg/ml) to inhibit cell proliferation. The wound healing area was imaged using an EVOS FL Auto2 microscope (ThermoFisher) and measured using Fiji ImageJ software.

### Transwell migration and invasion assays

Transwell chambers with 8-µm pore sized membranes (353097, Corning) and Matrigel invasion chambers with 8-µm pore sized membranes (354483, Corning) were used for migration and invasion assays, respectively. Briefly, 5x10^4^ cells in DMEM without FBS were added to the upper chamber of the inserts, while DMEM supplemented with 10% FBS was added to the lower compartment. Cells were incubated for 16 hours. Cells on the upper side of the membrane were gently scraped and washed off with PBS before the inserts were immersed in ice-cold methanol for 20 minutes at RT to fix the cells on the lower side of the membrane. The cells were then stained with 0.1% Crystal Violet solution for 20 minutes at RT, washed three times with distilled water, and then imaged using a Leica DM IL LED microscope. Cell numbers were quantified in 5 random fields using Fiji ImageJ software.

### Statistics

Statistical analyses were performed using GraphPad Prism v.10 (GraphPad Software). Tests used, multi-comparison correction methods, and significance cutoff were indicated in the figure legends for each quantification. The calculated *P*-values were shown in each graph.

---

**Figure EV1.**
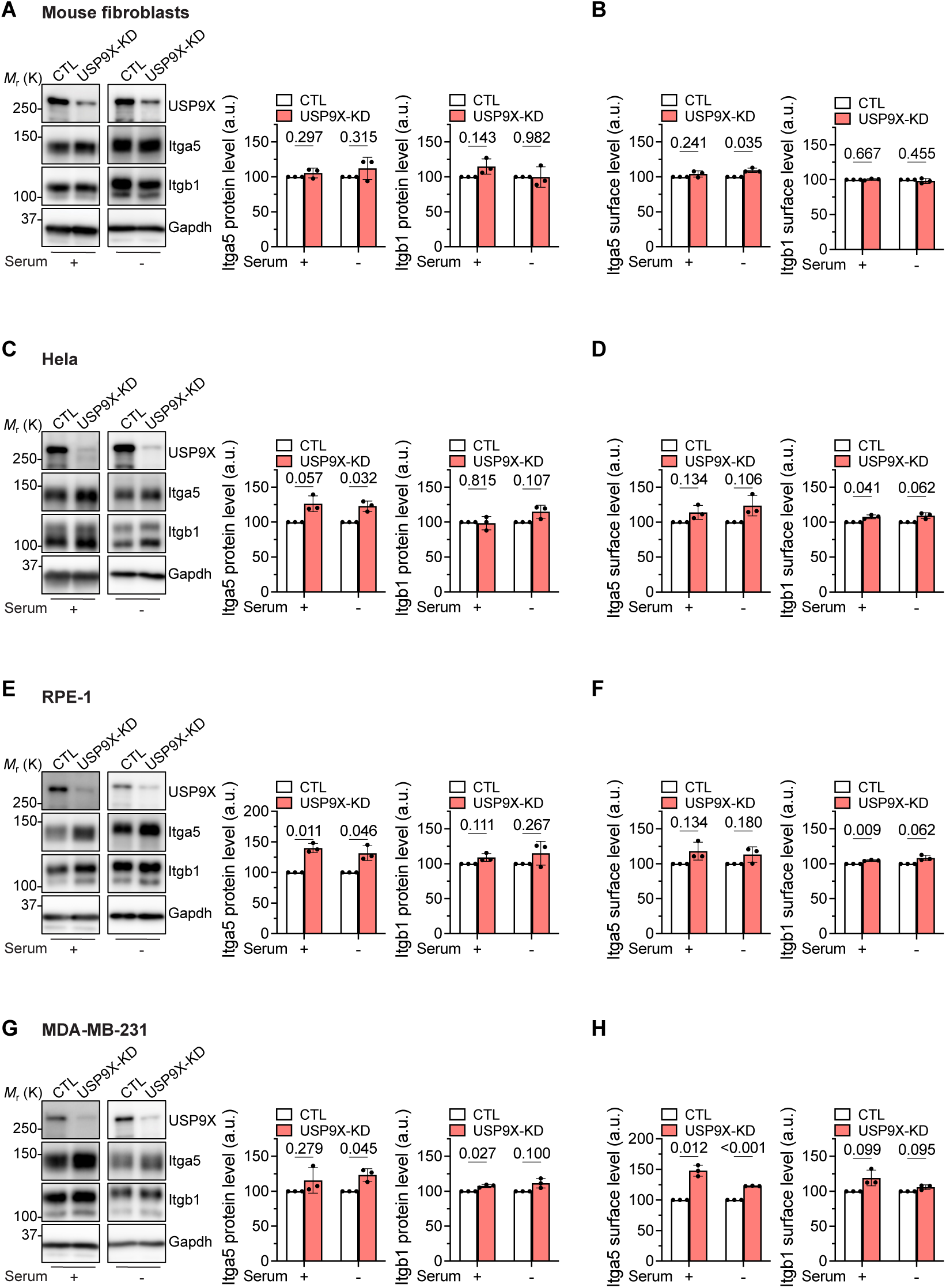
The effect of USP9X-KD on integrin levels. **(A-H)** WB and densitometric quantification of Itga5 and Itgb1 protein levels (**A**, **C**, **E**, **G**) and flow cytometry analysis of Itga5 and Itgb1 surface levels (**B**, **D**, **F**, **H**) in mouse fibroblasts (**A**, **B**), Hela (**C**, **D**), RPE-1 (**E**, **F**) and MDA-MB-231 cells (**G**, **H**) treated with control non-targeting siRNA (CTL) or siRNAs targeting USP9X (USP9X-KD). Cells were cultured overnight in DMEM with 10% serum or serum-replacement medium. Gapdh served as a loading control. Statistical analysis was carried out by paired t-test. Data are shown as Mean±SD, n=3 independent experiments.

**Figure EV2.**
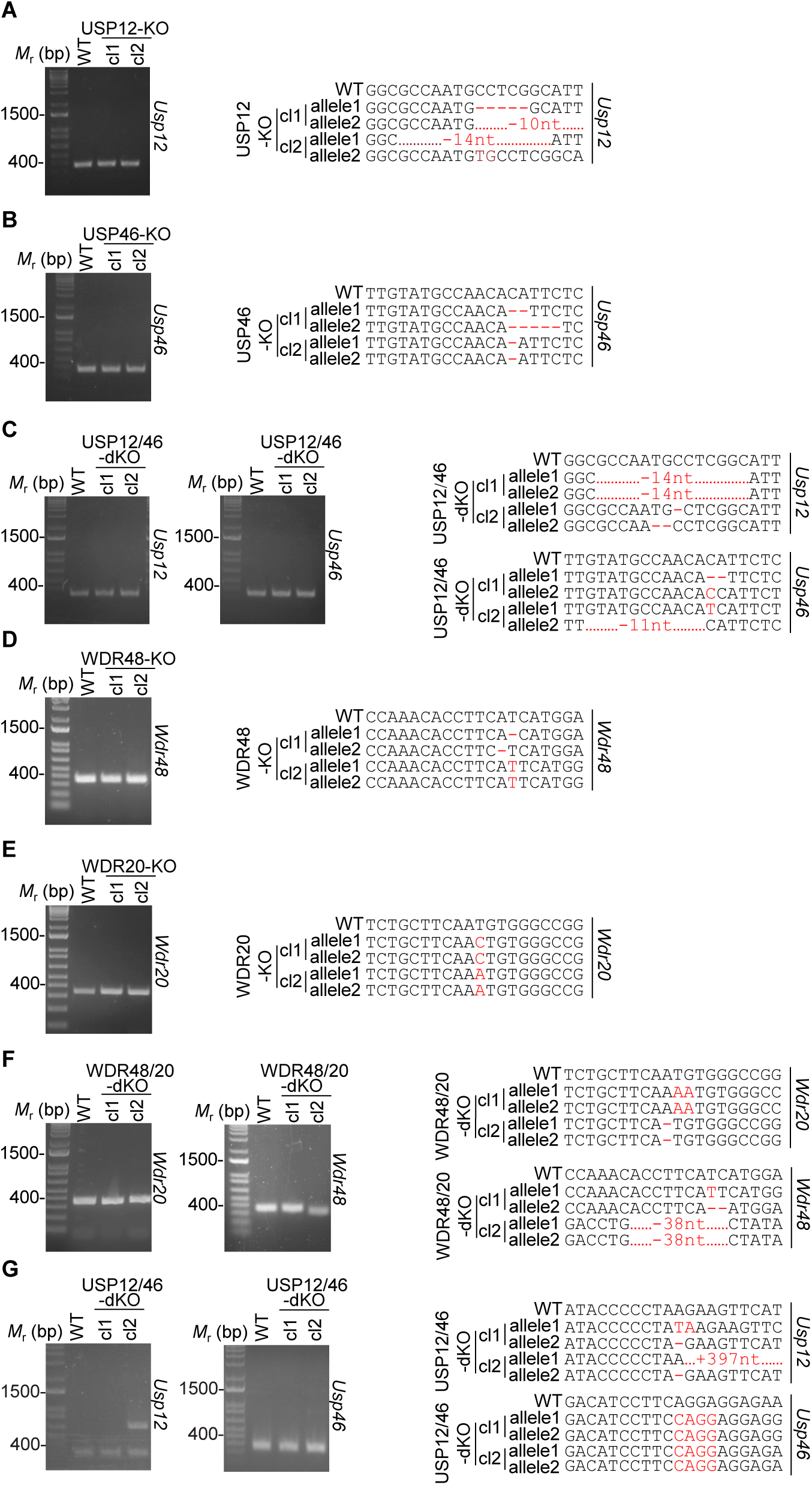
Validation of KO clones. **(A-G)** Agarose gel electrophoresis images show PCR amplification products from the genomic region containing the indicated Cas9 targeting sites of the indicated genes in the parental WT and two independent mouse fibroblast clones (**A-F**) and MDA-MB-231 cell clones (**G**). PCR products were sequenced, analyzed with the Synthego Inference of CRISPR Edits (ICE) analysis tool (Hsiau *et al*, 2018) and the corresponding alignments are shown on the right panel.

**Figure EV3.**
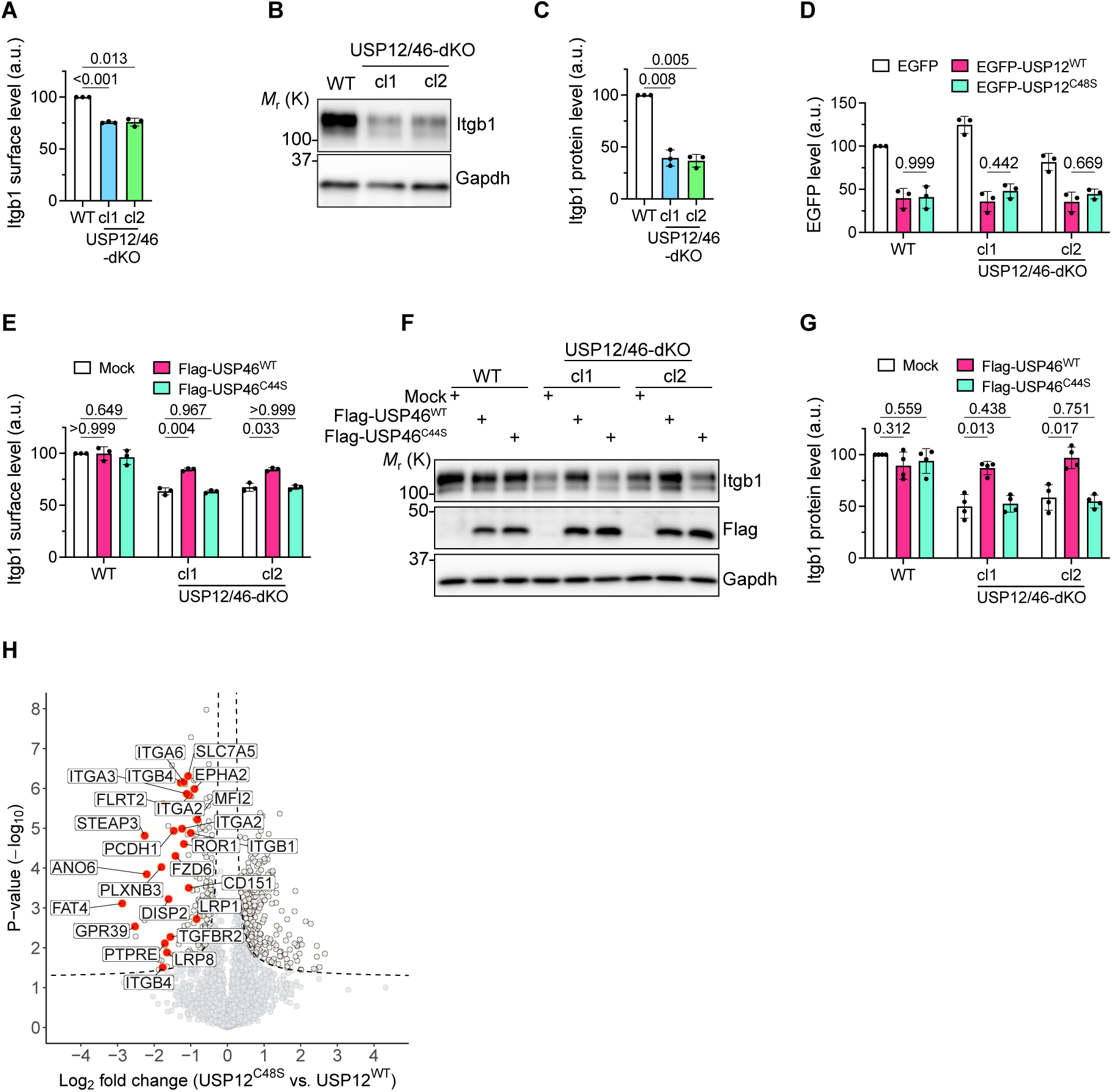
The USP12/46-WDRs complex stabilizes Itgb1 protein levels. **(A-C)** Itgb1 surface levels determined by flow cytometry (**A**) and Itgb1 protein levels in cell lysates determined by WB (**B**) with densitometric quantification (**C**) in WT and USP12/46-dKO MDA-MB-231 cells. Gapdh served as a loading control. Statistical analysis was carried out by RM one-way ANOVA with Dunnett’s multiple comparison test comparing with WT. Data are shown as Mean±SD, n=3 independent experiments. **(D)** EGFP fluorescence intensities in WT and USP12/46-dKO fibroblasts stably expressing EGFP, EGFP-USP12^WT^ or EGFP-USP12^C48S^ determined by flow cytometry. Statistical analysis was carried out by ordinary two-way ANOVA with Šidák’s multiple comparison test. Data are shown as Mean±SD, n=3 independent experiments. **(E-G)** Itgb1 surface levels determined by flow cytometry (**E**) and Itgb1 protein levels in cell lysates determined by WB (**F**) with densitometric quantification (**G**) in WT and USP12/46-dKO fibroblasts stably expressing FLAG-USP46^WT^ or FLAG-USP46^C44S^. Mock transduced cells (Mock) served as control. Gapdh served as a loading control. Statistical analysis was carried out by RM two-way ANOVA with Dunnett’s multiple comparison test. Data are shown as Mean±SD. **E**, n=3; **F, G** n=4 independent experiments. **(H)** Volcano plot of the cell surface proteome of USP12/46-dKO MDA-MB-231 cells expressing EGFP-USP12^C48S^ versus EGFP-USP12^WT^ identified by label-free MS. *P*-values are determined using two-sided permuted t-test with 250 randomizations. The black dashed line indicates the significance cutoff (FDR:0.05, S0:0.1) estimated by the Perseus software. n=4 biological replicates. Arbitrarily selected cell surface receptors are highlighted in red.

**Figure EV4.**
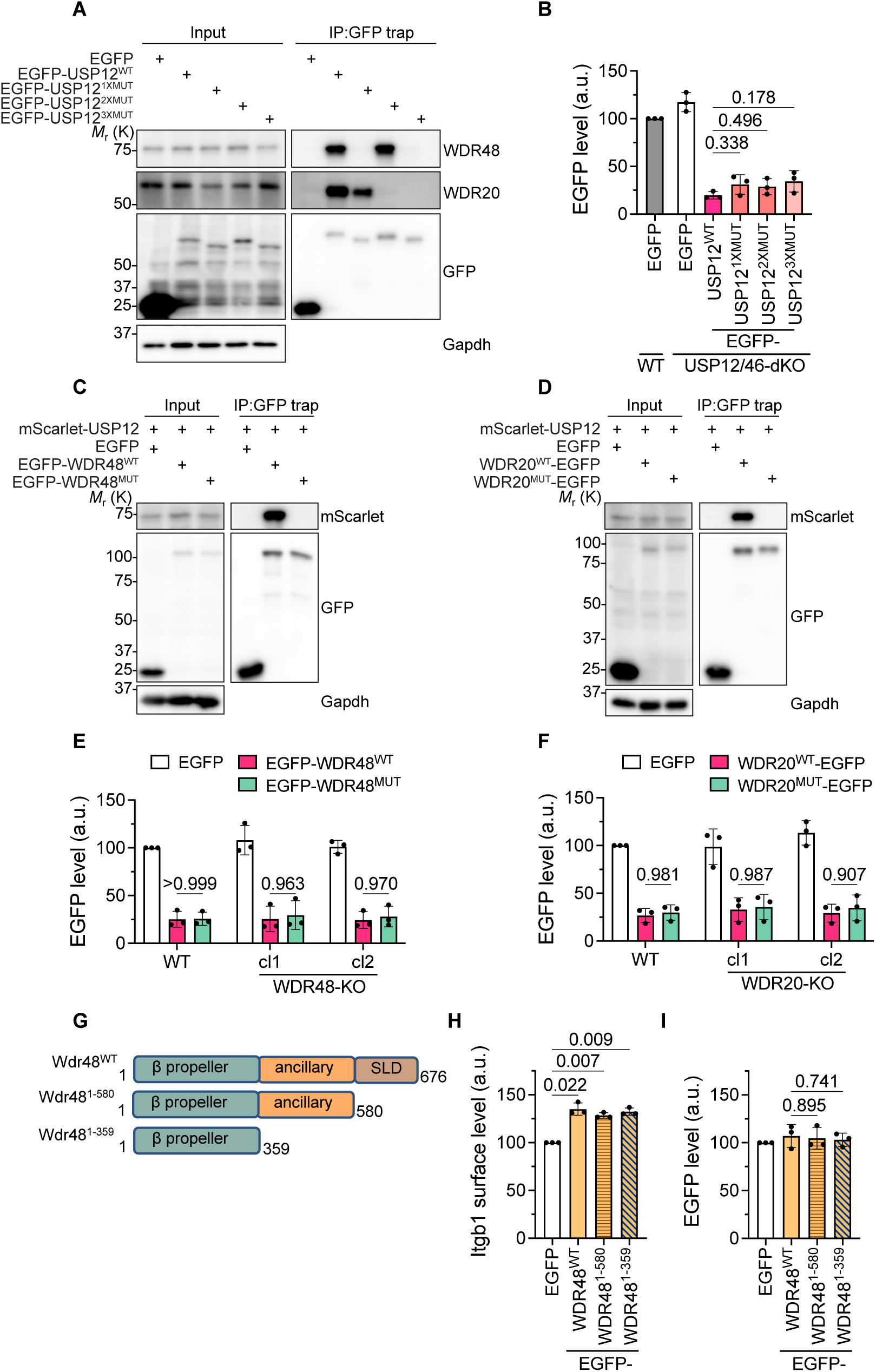
Characterization of binding-deficient USP12, WDR48 and WDR20 mutants. **(A)** GFP immunoprecipitation (GFP IP) from USP12/46-dKO fibroblasts transiently expressing EGFP, EGFP-USP12^WT^, EGFP-USP12^1XMUT^, EGFP-USP12^2XMUT^ or EGFP-USP12^3XMUT^ analyzed by WB for indicated proteins. Gapdh served as a loading control. Representative images from 3 independent experiments are shown. **(B)** EGFP fluorescence in WT and USP12/46-dKO fibroblasts transiently expressing EGFP, EGFP-USP12^WT^, EGFP-USP12^1XMUT^, EGFP-USP12^2XMUT^ or EGFP-USP12^3XMUT^ determined by flow cytometry. Statistical analysis was carried out by ordinary one-way ANOVA with Dunnett’s multiple comparison test. Data are shown as Mean±SD, n=3 independent experiments. **(C)** GFP IP from USP12/46-dKO fibroblasts stably expressing mScarlet-USP12 and transiently expressing EGFP, EGFP-WDR48^WT^ or EGFP-WDR48^MUT^ analyzed by WB for indicated proteins. Gapdh served as a loading control. Representative images from 3 independent experiments are shown. **(D)** GFP IP from USP12/46-dKO fibroblasts stably expressing mScarlet-USP12 and transiently expressing EGFP, WDR20^WT^-EGFP or WDR20^MUT^-EGFP analyzed by WB for indicated proteins. Gapdh served as a loading control. Representative images from 3 independent experiments are shown. **(E)** EGFP fluorescence in WT and WDR48-KO fibroblasts transiently expressing EGFP, EGFP-WDR48^WT^ or EGFP-WDR48^MUT^ determined by flow cytometry. Statistical analysis was carried out by ordinary two-way ANOVA with Šidák’s multiple comparison test. Data are shown as Mean±SD, n=3 independent experiments. **(F)** EGFP fluorescence in WT and WDR20-KO fibroblasts transiently expressing EGFP, WDR20^WT^-EGFP or WDR20^MUT^-EGFP determined by flow cytometry. Statistical analysis was carried out by ordinary two-way ANOVA with Šidák’s multiple comparison test. Data are shown as Mean±SD, n=3 independent experiments. **(G)** Domain organization of the WT WDR48 and WDR48 domain-deletion mutants. **(H, I)** Itgb1 surface levels (**H**) and EGFP fluorescence (**I**) in WDR48-KO fibroblasts stably expressing EGFP, EGFP-WDR48^WT^, EGFP-WDR48^1-580^ or EGFP-WDR48^1-359^ determined by flow cytometry. Statistical analysis was carried out by RM one-way ANOVA with Dunnett’s multiple comparison test. Data are shown as Mean±SD, n=3 independent experiments.

**Figure EV5.**
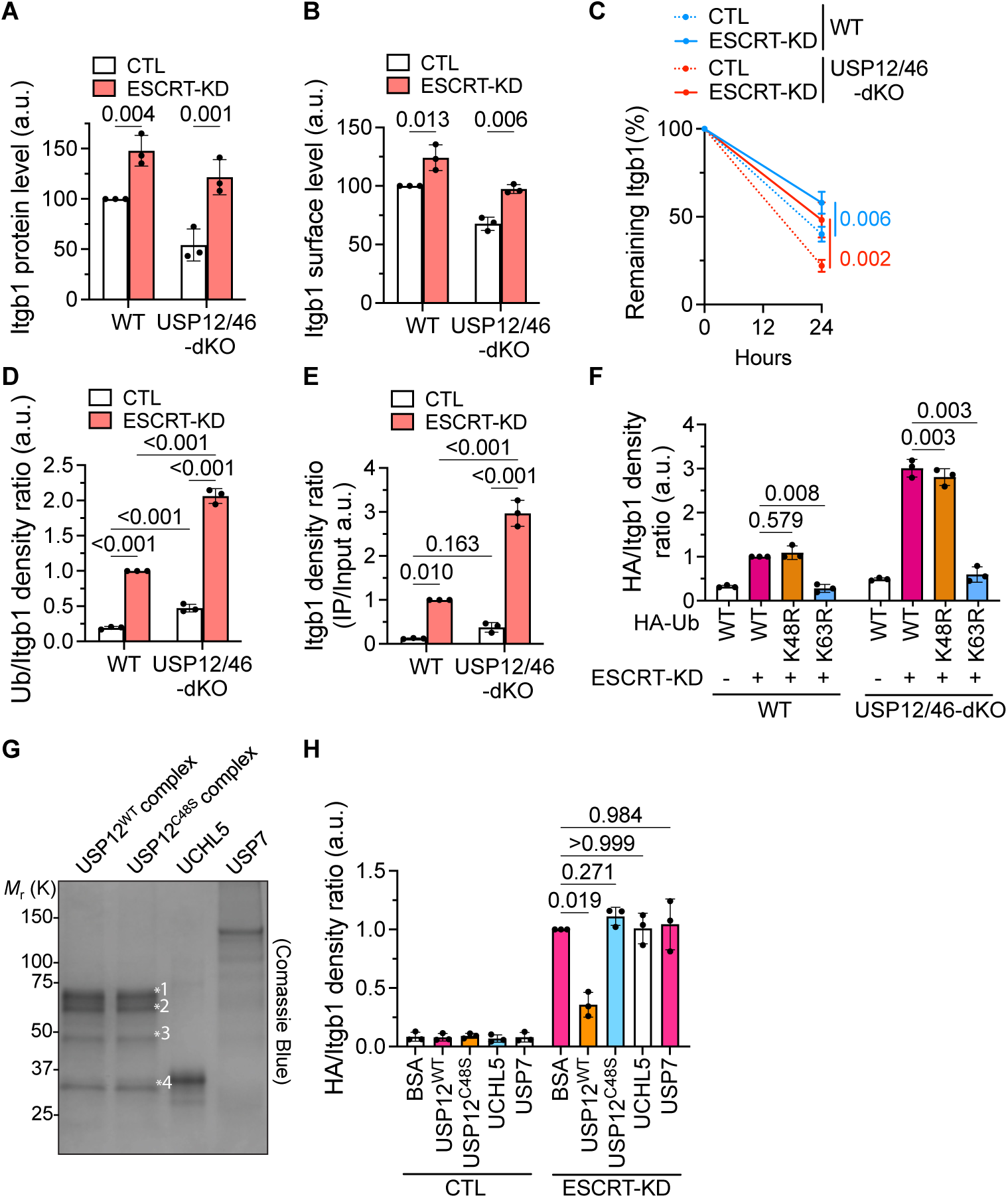
The ubiquitin-ESCRT pathway mediates Itgb1 degradation. **(A)** Densitometric quantification of Itgb1 proteins levels in Fig. 5A. Statistical analysis was carried out by RM two-way ANOVA with Šidák’s multiple comparison test. Data are shown as Mean±SD, n=3 independent experiments. **(B)** Itgb1 surface levels in WT and USP12/46-dKO fibroblasts treated with or without ESCRT-KD siRNAs determined by flow cytometry. Statistical analysis was carried out by RM two-way ANOVA with Šidák’s multiple comparison test. Data are shown as Mean±SD, n=3 independent experiments. **(C)** Surface Itgb1 degradation kinetics in WT and USP12/46-dKO fibroblasts treated with or without ESCRT-KD siRNAs measured by capture-ELISA. Statistical analysis was carried out by RM two-way ANOVA with Šidák’s multiple comparison test. Data are shown as Mean±SD, n=3 independent experiments. **(D)** Quantification of Itgb1 ubiquitination levels in Fig. 5B. The intensity of the Ub signals was normalized to the intensity of the IP-ed Itgb1 signals. Statistical analysis was carried out by RM two-way ANOVA with Šidák’s multiple comparison test. Data are shown as Mean±SD, n=3 independent experiments. **(E)** Quantification of Itgb1 ubiquitination levels in Fig. 5C. The intensity of the IP-ed Itgb1 signals was normalized to the intensity of the input Itgb1 signals. Statistical analysis was carried out by RM two-way ANOVA with Šidák’s multiple comparison test. Data are shown as Mean±SD, n=3 independent experiments. **(F)** Quantification of Itgb1 ubiquitination levels in Fig. 5E. The intensity of the HA signals was normalized to the intensity of the IP-ed Itgb1 signals. Statistical analysis was carried out by RM two-way ANOVA with Dunnett’s multiple comparison test. Data are shown as Mean±SD, n=3 independent experiments. **(G)** Coomassie-blue staining of recombinant proteins used in the *in vitro* de-ubiquitination assay. MS was used to determine the identity of the protein bands indicated in the DUB complexes. Band1, WDR48; band2, WDR20; band3, a cleaved WDR20; band4, USP12. **(H)** Quantification of Itgb1 ubiquitination levels in Fig. 5F. The intensity of the Ub signals was normalized to the intensity of the IP-ed Itgb1 signals. Statistical analysis was carried out by RM two-way ANOVA with Dunnett’s multiple comparison test. Data are shown as Mean±SD, n=3 independent experiments.

**Figure EV6.**
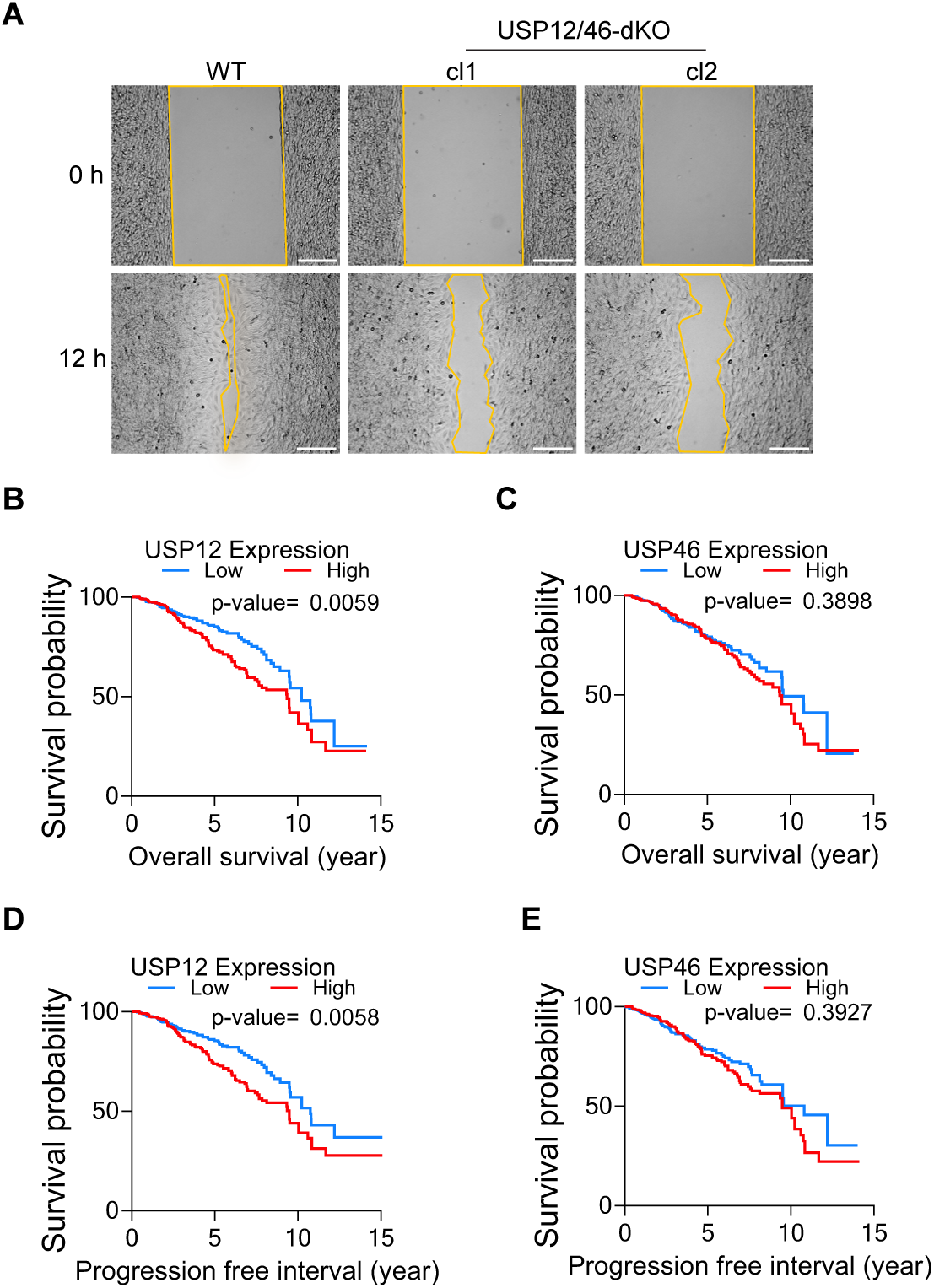
USP12 and USP46 are not favorable for prognosis in cancer patients. **(A)** Representative images of the *in vitro* wound healing assay showing WT and USP12/46-dKO fibroblasts migrating on FN-coated 2D surfaces at 0 and 12 hours. Lines mark the leading edge of cell migration towards the wound. Scale bar, 200 µm. **(B-E)** Kaplan Meier-plot of the overall survival (**B, D**) and progression free interval (**C, E**) of breast cancer patients with high (red line) or low (blue line) gene expression *USP12* (**B, C**) or *USP46* (**D, E**) levels. The GDC TCGA dataset obtained from the UCSC Xena project (Goldman *et al*., 2020) was used. Two-group risk model with cut-off at the median was applied. *P*-values were calculated by Log-rank test.

## References

1. Bottcher RT, Stremmel C, Meves A, Meyer H, Widmaier M, Tseng HY, Fassler R (2012) Sorting nexin 17 prevents lysosomal degradation of beta1 integrins by binding to the beta1-integrin tail. Nat Cell Biol 14: 584–592

2. Branon TC, Bosch JA, Sanchez AD, Udeshi ND, Svinkina T, Carr SA, Feldman JL, Perrimon N, Ting AY (2018) Efficient proximity labeling in living cells and organisms with TurboID. Nat Biotechnol 36: 880–887

3. Calderwood DA, Campbell ID, Critchley DR (2013) Talins and kindlins: partners in integrin-mediated adhesion. Nat Rev Mol Cell Biol 14: 503–517

4. Clague MJ, Liu H, Urbe S (2012) Governance of endocytic trafficking and signaling by reversible ubiquitylation. Dev Cell 23: 457–467

5. Dharadhar S, Clerici M, van Dijk WJ, Fish A, Sixma TK (2016) A conserved two-step binding for the UAF1 regulator to the USP12 deubiquitinating enzyme. J Struct Biol 196: 437–447

6. Dozynkiewicz MA, Jamieson NB, Macpherson I, Grindlay J, van den Berghe PV, von Thun A, Morton JP, Gourley C, Timpson P, Nixon C et al (2012) Rab25 and CLIC3 collaborate to promote integrin recycling from late endosomes/lysosomes and drive cancer progression. Dev Cell 22: 131–145

7. Fitzpatrick P, Shattil SJ, Ablooglu AJ (2014) C-terminal COOH of integrin beta1 is necessary for beta1 association with the kindlin-2 adapter protein. J Biol Chem 289: 11183–11193

8. Goldman MJ, Craft B, Hastie M, Repecka K, McDade F, Kamath A, Banerjee A, Luo Y, Rogers D, Brooks AN et al (2020) Visualizing and interpreting cancer genomics data via the Xena platform. Nat Biotechnol 38: 675–678

9. Groza T, Gomez FL, Mashhadi HH, Munoz-Fuentes V, Gunes O, Wilson R, Cacheiro P, Frost A, Keskivali-Bond P, Vardal B et al (2023) The International Mouse Phenotyping Consortium: comprehensive knockout phenotyping underpinning the study of human disease. Nucleic Acids Res 51: D1038–D1045

10. Hanson PI, Cashikar A (2012) Multivesicular Body Morphogenesis. Annual Review of Cell and Developmental Biology 28: 337–362

11. Horton ER, Byron A, Askari JA, Ng DHJ, Millon-Fremillon A, Robertson J, Koper EJ, Paul NR, Warwood S, Knight D et al (2015) Definition of a consensus integrin adhesome and its dynamics during adhesion complex assembly and disassembly. Nat Cell Biol 17: 1577–1587

12. Hsiau T, Conant D, Rossi N, Maures T, Waite K, Yang J, Joshi S, Kelso R, Holden K, Enzmann BL, Stoner R, 2018. Inference of CRISPR Edits from Sanger Trace Data. Cold Spring Harbor Laboratory. Huotari J, Helenius A (2011) Endosome maturation. EMBO J 30: 3481–3500

13. Hynes RO (2002) Integrins: bidirectional, allosteric signaling machines. Cell 110: 673–687

14. Kharitidi D, Apaja PM, Manteghi S, Suzuki K, Malitskaya E, Roldan A, Gingras MC, Takagi J, Lukacs GL, Pause A (2015) Interplay of Endosomal pH and Ligand Occupancy in Integrin alpha5beta1 Ubiquitination, Endocytic Sorting, and Cell Migration. Cell Rep 13: 599–609

15. Komander D, Clague MJ, Urbé S (2009) Breaking the chains: structure and function of the deubiquitinases. Nature Reviews Molecular Cell Biology 10: 550–563

16. Kuo J-C, Han X, Hsiao C-T, Yates Iii JR, Waterman CM (2011) Analysis of the myosin-II-responsive focal adhesion proteome reveals a role for β-Pix in negative regulation of focal adhesion maturation. Nature Cell Biology 13: 383–393

17. Li H, Deng Y, Sun K, Yang H, Liu J, Wang M, Zhang Z, Lin J, Wu C, Wei Z, Yu C (2017) Structural basis of kindlin-mediated integrin recognition and activation. Proc Natl Acad Sci U S A 114: 9349–9354

18. Li H, Lim KS, Kim H, Hinds TR, Jo U, Mao H, Weller CE, Sun J, Chatterjee C, D’Andrea AD, Zheng N (2016) Allosteric Activation of Ubiquitin-Specific Proteases by beta-Propeller Proteins UAF1 and WDR20. Mol Cell 63: 249–260

19. Li W, Xu H, Xiao T, Cong L, Love MI, Zhang F, Irizarry RA, Liu JS, Brown M, Liu XS (2014) MAGeCK enables robust identification of essential genes from genome-scale CRISPR/Cas9 knockout screens. Genome Biology 15

20. Lim KL, Chew KC, Tan JM, Wang C, Chung KK, Zhang Y, Tanaka Y, Smith W, Engelender S, Ross CA et al (2005) Parkin mediates nonclassical, proteasomal-independent ubiquitination of synphilin-1: implications for Lewy body formation. J Neurosci 25: 2002–2009

21. Lobert VH, Brech A, Pedersen NM, Wesche J, Oppelt A, Malerod L, Stenmark H (2010) Ubiquitination of alpha 5 beta 1 integrin controls fibroblast migration through lysosomal degradation of fibronectin-integrin complexes. Dev Cell 19: 148–159

22. McNally KE, Faulkner R, Steinberg F, Gallon M, Ghai R, Pim D, Langton P, Pearson N, Danson CM, Nagele H et al (2017) Retriever is a multiprotein complex for retromer-independent endosomal cargo recycling. Nat Cell Biol 19: 1214–1225

23. Melamed S, Zaffryar-Eilot S, Nadjar-Boger E, Aviram R, Zhao H, Yaseen-Badarne W, Kalev-Altman R, Sela-Donenfeld D, Lewinson O, Astrof S et al (2023) Initiation of fibronectin fibrillogenesis is an enzyme-dependent process. Cell Reports 42: 112473

24. Meng Y, Hong C, Yang S, Qin Z, Yang L, Huang Y (2023) Roles of USP9X in cellular functions and tumorigenesis (Review). Oncology Letters 26

25. Miranda M, Sorkin A (2007) Regulation of receptors and transporters by ubiquitination: new insights into surprisingly similar mechanisms. Mol Interv 7: 157–167

26. Moreno-Layseca P, Icha J, Hamidi H, Ivaska J (2019) Integrin trafficking in cells and tissues. Nat Cell Biol 21: 122–132

27. Moser M, Legate KR, Zent R, Fassler R (2009) The tail of integrins, talin, and kindlins. Science 324: 895–899

28. Nathan JA, Kim HT, Ting L, Gygi SP, Goldberg AL (2013) Why do cellular proteins linked to K63-polyubiquitin chains not associate with proteasomes? EMBO J 32: 552–565

29. Niu K, Shi Y, Lv Q, Wang Y, Chen J, Zhang W, Feng K, Zhang Y (2023) Spotlights on ubiquitin-specific protease 12 (USP12) in diseases: from multifaceted roles to pathophysiological mechanisms. J Transl Med 21: 665

30. Paulmann C, Spallek R, Karpiuk O, Heider M, Schaffer I, Zecha J, Klaeger S, Walzik M, Ollinger R, Engleitner T et al (2022) The OTUD6B-LIN28B-MYC axis determines the proliferative state in multiple myeloma. EMBO J 41: e110871

31. Saftig P, Klumperman J (2009) Lysosome biogenesis and lysosomal membrane proteins: trafficking meets function. Nat Rev Mol Cell Biol 10: 623–635

32. Schiller HB, Friedel CC, Boulegue C, Fässler R (2011) Quantitative proteomics of the integrin adhesome show a myosin II-dependent recruitment of LIM domain proteins. EMBO reports 12: 259–266

33. Steinberg F, Heesom KJ, Bass MD, Cullen PJ (2012) SNX17 protects integrins from degradation by sorting between lysosomal and recycling pathways. J Cell Biol 197: 219–230

34. Strickland M, Watanabe S, Bonn SM, Camara CM, Starich MR, Fushman D, Carter CA, Tjandra N (2022) Tsg101/ESCRT-I recruitment regulated by the dual binding modes of K63-linked diubiquitin. Structure 30: 289–299 e286

35. Swatek KN, Komander D (2016) Ubiquitin modifications. Cell Res 26: 399–422

36. Uhlen M, Fagerberg L, Hallstrom BM, Lindskog C, Oksvold P, Mardinoglu A, Sivertsson A, Kampf C, Sjostedt E, Asplund A et al (2015) Proteomics. Tissue-based map of the human proteome. Science 347: 1260419

37. Wang R, Wang J, Hassan A, Lee CH, Xie XS, Li X (2021) Molecular basis of V-ATPase inhibition by bafilomycin A1. Nat Commun 12: 1782

38. Xu G, Paige JS, Jaffrey SR (2010) Global analysis of lysine ubiquitination by ubiquitin remnant immunoaffinity profiling. Nat Biotechnol 28: 868–873

39. Yin J, Schoeffler AJ, Wickliffe K, Newton K, Starovasnik MA, Dueber EC, Harris SF (2015) Structural Insights into WD-Repeat 48 Activation of Ubiquitin-Specific Protease 46. Structure 23: 2043–2054

40. Zhu H, Zhang T, Wang F, Yang J, Ding J (2019) Structural insights into the activation of USP46 by WDR48 and WDR20. Cell Discov 5: 34

## References

41. Aretz J, Aziz M, Strohmeyer N, Sattler M, Fassler R (2023) Talin and kindlin use integrin tail allostery and direct binding to activate integrins. Nat Struct Mol Biol 30: 1913–1924

42. Azimifar SB, Bottcher RT, Zanivan S, Grashoff C, Kruger M, Legate KR, Mann M, Fassler R (2012) Induction of membrane circular dorsal ruffles requires co-signalling of integrin-ILK-complex and EGF receptor. J Cell Sci 125: 435–448

43. Benito-Jardon M, Klapproth S, Gimeno LI, Petzold T, Bharadwaj M, Muller DJ, Zuchtriegel G, Reichel CA, Costell M (2017) The fibronectin synergy site re-enforces cell adhesion and mediates a crosstalk between integrin classes. Elife 6

44. Böttcher RT, Stremmel C, Meves A, Meyer H, Widmaier M, Tseng HY, Fässler R (2012) Sorting nexin 17 prevents lysosomal degradation of beta1 integrins by binding to the beta1-integrin tail. Nat Cell Biol 14: 584–592

45. Chen NP, Aretz J, Fassler R (2022) CDK1-cyclin-B1-induced kindlin degradation drives focal adhesion disassembly at mitotic entry. Nat Cell Biol 24: 723–736

46. Cox J, Mann M (2008) MaxQuant enables high peptide identification rates, individualized p.p.b.-range mass accuracies and proteome-wide protein quantification. Nat Biotechnol 26: 1367–1372

47. Galaxy C (2022) The Galaxy platform for accessible, reproducible and collaborative biomedical analyses: 2022 update. Nucleic Acids Res 50: W345–W351

48. Li H, Lim KS, Kim H, Hinds TR, Jo U, Mao H, Weller CE, Sun J, Chatterjee C, D’Andrea AD et al (2016) Allosteric Activation of Ubiquitin-Specific Proteases by beta-Propeller Proteins UAF1 and WDR20. Mol Cell 63: 249–260

49. Lobert VH, Brech A, Pedersen NM, Wesche J, Oppelt A, Malerod L, Stenmark H (2010) Ubiquitination of alpha 5 beta 1 integrin controls fibroblast migration through lysosomal degradation of fibronectin-integrin complexes. Dev Cell 19: 148–159

50. Paulmann C, Spallek R, Karpiuk O, Heider M, Schaffer I, Zecha J, Klaeger S, Walzik M, Ollinger R, Engleitner T et al (2022) The OTUD6B-LIN28B-MYC axis determines the proliferative state in multiple myeloma. EMBO J 41: e110871

51. Ran FA, Hsu PD, Wright J, Agarwala V, Scott DA, Zhang F (2013) Genome engineering using the CRISPR-Cas9 system. Nat Protoc 8: 2281–2308

52. Sanjana NE, Shalem O, Zhang F (2014) Improved vectors and genome-wide libraries for CRISPR screening. Nature Methods 11: 783–784

53. Stauffer W, Sheng H, Lim HN (2018) EzColocalization: An ImageJ plugin for visualizing and measuring colocalization in cells and organisms. Sci Rep 8: 15764

54. Theodosiou M, Widmaier M, Bottcher RT, Rognoni E, Veelders M, Bharadwaj M, Lambacher A, Austen K, Muller DJ, Zent R et al (2016) Kindlin-2 cooperates with talin to activate integrins and induces cell spreading by directly binding paxillin. Elife 5: e10130

